# Chromosome-specific drift under stabilizing selection generates polygenic barriers to sex chromosome turnover

**DOI:** 10.1101/2024.09.03.611135

**Authors:** Pavitra Muralidhar

## Abstract

Sex chromosome systems show frequent evolutionary transitions in some clades, but long-term stability in others. Previous explanations of this stasis rely on evolutionary dynamics peculiar to sex chromosomes, such as the accumulation of deleterious mutations on the sex-specific chromosome or sexually antagonistic mutations on either sex chromosome. Here, I show that stabilizing selection on quantitative traits promotes stability of sex chromosome systems. The reason is that stabilizing selection, while keeping the value of the trait near its optimum, allows individual chromosomes’ contributions to the trait to drift, and this chromosome-specific drift reduces the fitness of the novel sexual genotypes necessarily produced during sex chromosome turnover. Given the ubiquity of stabilizing selection on quantitative traits, chromosome-specific drift could play a pivotal role in preventing the turnover of sex chromosome systems across multiple stages of their evolution and can explain key patterns in the phylogenetic distribution of sex-determining systems.

---

The theory rests on three well-supported assumptions. First, it requires that some quantitative traits—continuously distributed and controlled by many genes throughout the genome, including on the sex chromosomes—are under stabilizing selection, that is, selection for an ‘optimal’ trait value. Genomic research over the past few decades has established that most heritable quantitative traits have a polygenic basis. Empirical evidence and theoretical expectation further point to many of these traits being under stabilizing selection (1–4).

Second, the theory requires that the genetic variants controlling these traits have different— though possibly correlated—effects in males and females. Many quantitative traits have a shared, but not identical, genetic architecture in males and females, permitting sexual dimorphism in these phenotypes (2, 5–7), and empirical studies across taxa have estimated strong, but not perfect, intersex correlations of phenotypic effects (6, 8–10).

Third, the theory requires that the sex chromosomes do not recombine along at least a portion of their length in the heterogametic sex. Recombination shutdown between the X and Y (or Z and W) chromosomes is a common property of heterogametic systems, including young systems with little divergence between the X and Y; the size of the non-recombining region varies widely across systems and often increases over time (11–14). The mechanisms underlying recombination shutdown are diverse, and the theory presented here does not rely on any particular mechanism (12, 15–18)

Note that, in this model, the fitness effects of individual alleles are not specified exogenously. This is in contradistinction to classical theories of sex chromosome turnover, which typically designate alleles to be beneficial in one sex and detrimental in the other, or always beneficial or always deleterious (19–23). Instead, in the model considered here, alleles simply contribute—in varying directions and degrees—to polygenic phenotypes. Their fitness effects, and hence the population genetic dynamics at their loci, emerge endogenously via selection on these phenotypes (4–5, 24–26). This approach flows from our modern understanding of the genetic architecture of complex traits, and represents a new framework, grounded in well-established principles of quantitative genetic theory, for modeling the evolution of sex-determining systems.

Under the three assumptions set out above, the genetic dynamics of a quantitative trait can be studied precisely. Stabilizing selection causes the average additive genetic value of the trait to evolve rapidly to its optimum, where it is then constrained by selection to remain (24). Crucially, however, while the genome-wide genetic value is constrained to equal the trait optimum, this is not true for subregions of the genome such as individual chromosomes. Instead, their average genetic values can drift upwards or downwards, subject only to the constraint that together their sum must equal the optimum. Thus, for example, chromosome 1’s average genetic value for the trait can increase if this is compensated for by a decrease in chromosome 2’s average genetic value. The resulting process of constrained drift can be modelled as a system of coupled stochastic differential equations (27, Appendix Section S1), the solution of which reveals that individual chromosomes’ genetic values can diverge rapidly even under strong stabilizing selection on their combined product (Fig. 1).

**Fig. 1.**
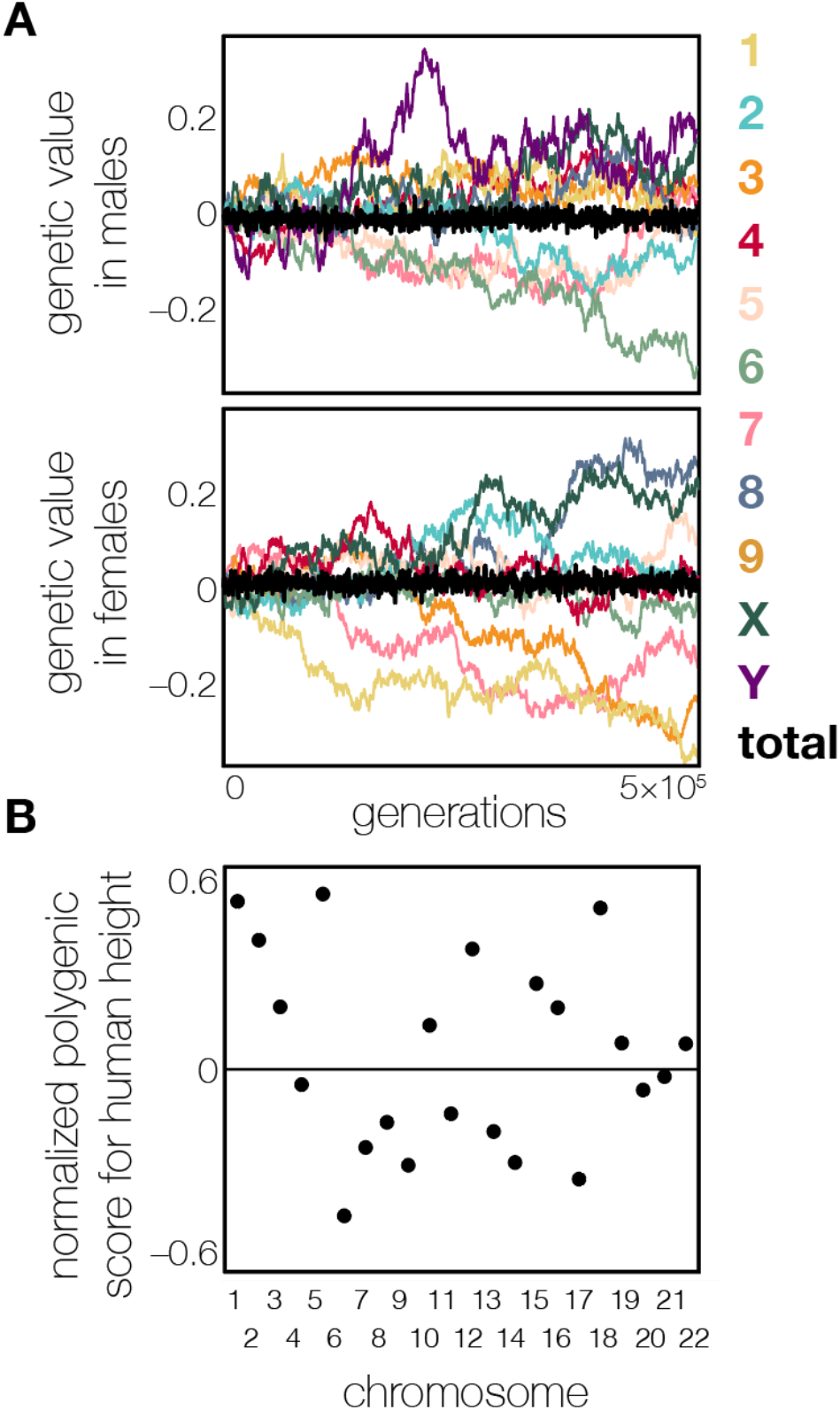
Chromosome-specific drift of genetic values for a trait under stabilizing selection. **(A)** Male and female genetic values of 10 chromosomes in a simulation of a polygenic trait under stabilizing selection around an optimum of 0. The genome-wide mean value remains close to 0 in both sexes (black trajectories), but the values of individual chromosomes vary increasingly over time (colored trajectories). Simulation details in Methods. **(B)** Variance across human autosomes in their average polygenic scores (PGSs) for height, a quantitative trait under stabilizing selection (2). Chromosome-specific PGSs were calculated from allele frequencies and GWAS effect sizes reported by Yengo et al. (65) and normalized by the average genome-wide value. Because the PGSs depend only on currently segregating genetic polymorphisms, they likely underestimate the true variation across chromosomes in their average genetic values for height.

Chromosome-specific drift under stabilizing selection has important implications for evolutionary transitions between sex chromosome systems. All such transitions—whether or not they recruit a new linkage group in sex determination, and whether or not they switch the system of heterogamety—generate novel sexual genotypes (21); that is, male or female genotypes that were not present in the initial sex chromosome system (Fig. 2). These novel sexual genotypes disrupt the dynamic cross-chromosome balancing act that had, under chromosome-specific drift, maintained quantitative traits at their optimal values under the original sex-determining system. The displacement from trait optima of novel sexual genotypes, and the concomitant reduction in their fitness, creates a barrier to the invasion of the new sex chromosome system. This barrier grows at a rate that depends on the genetic variance contributed by the non-recombining region of the sex chromosomes and, under empirically reasonable parameter regimes, can act as a substantial impediment to sex chromosome turnover. In the specification considered in Fig. 2, for example, the probability of a sex chromosome turnover decreases by approximately an order of magnitude across 500,000 generations (Fig. 2, S1, S2).

**Fig. 2.**
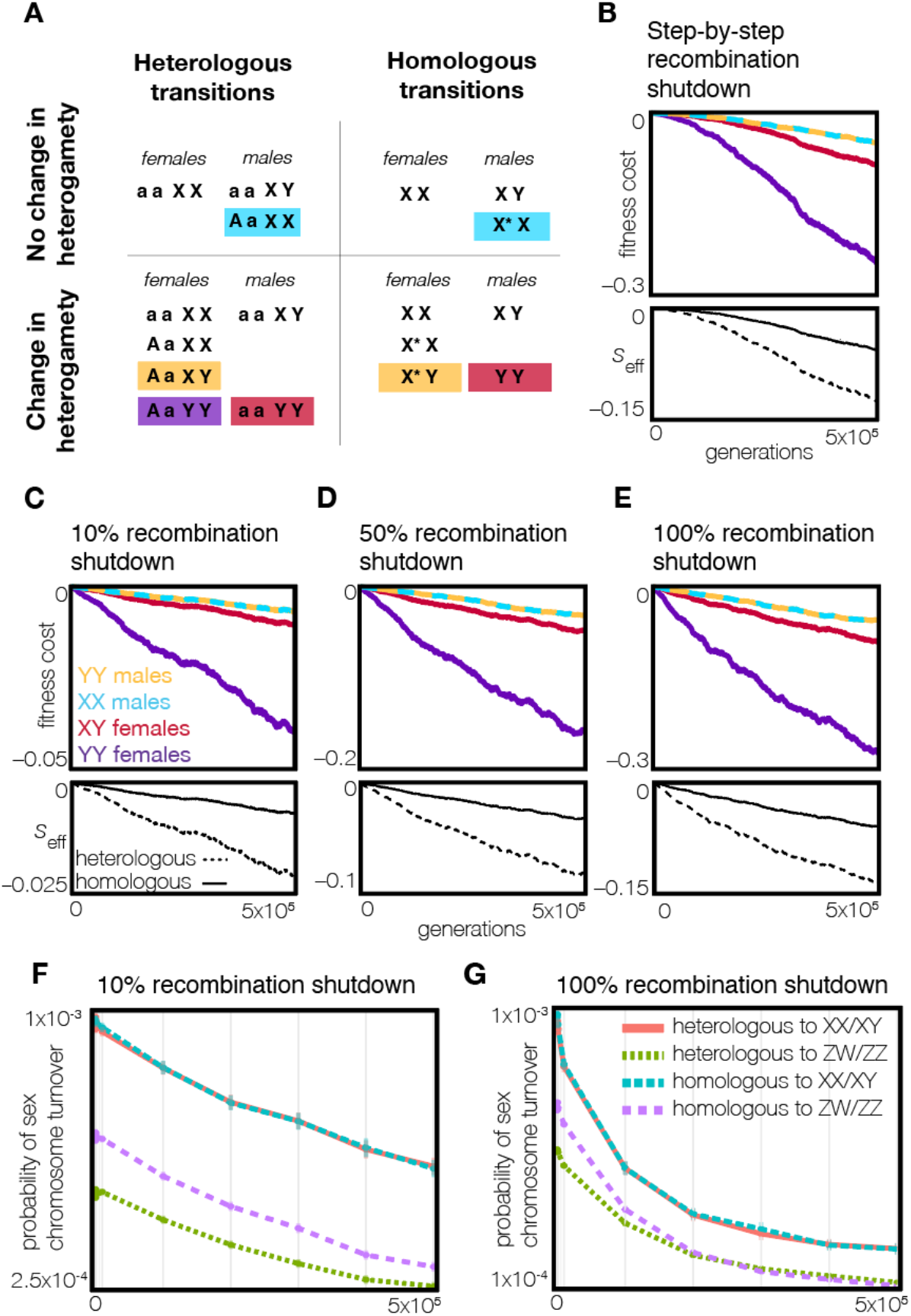
Chromosome-specific drift generates increasing barriers to sex chromosome turnover. **(A)** Sex chromosome turnovers always involve the creation of new sexual genotypes, whether they involve a new linkage group in sex determination (heterologous transitions; left) or not (homologous transitions; right), whether they change the heterogametic system (bottom) or not (top), and whether the new sex-determining mutations involved are dominant (here) or recessive (Fig. S12). **(B–E)** Chromosome-specific drift for sex-linked traits under stabilizing selection imposes fitness costs on new sexual genotypes, which increase with the age of the existing sex chromosome system. These costs scale with the extent of recombination suppression: (C) 10%, (D) 50%, or (E) complete X–Y recombination shutdown. (B) shows a scenario where recombination suppression expands by increments of 10% of the length of the chromosome every 2*N* generations. Eigenvalues of the corresponding dynamical systems (B–E) serve as an ‘effective selection coefficient’ *s*_eff_, providing a summary measure of the reduction in turnover likelihood, accounting for selection against the full set of sexual genotypes; this is especially useful for complex heterogametic transitions in which multiple intermediate sexual genotypes are involved in the transition (see Appendix S2 for more detail). **(F–G)** A sex chromosome turnover occurs when a new sex-determining mutation appears in a population and rises to fixation (25% frequency in the case of dominant sex-determining mutations). The probability of fixation of these new sex-determining mutations, and therefore a successful sex chromosomes turnover, declines as the X and Y diverge via chromosome-specific drift. Each point shows the probability of a sex chromosome turnover estimated from 10^7^ introductions of a single copy of a new sex-determining mutation. Simulations in B–E use population size *N* = 10,000 and per-locus mutation rate *μ* = 10^−5^; F–G use *N* = 1,000 and *μ* = 10^−6^, yielding equal rates of chromosome-specific drift (see Fig. S5, Methods). Because stochastic drift in smaller populations favors the invasion of new sex-determining mutations, F–G provide conservative estimates of turnover probabilities. All simulations model a single trait under stabilizing selection of strength similar to that estimated for human height (2).

For simplicity, I have described the theory of chromosome-specific drift above in terms of the genetic values of entirely non-recombining sex chromosomes. If only a subregion of the sex chromosomes has undergone recombination shutdown—as is common in the early stages of sex chromosome evolution—the fitness costs to novel sexual genotypes produced during a sex chromosome turnover will be proportional to the genetic variance contributed by the non-recombining region (Fig. 2C,D). Even if the non-recombining region constitutes a small portion of the sex chromosomes, the barrier to sex chromosome turnover caused by subregion drift can be quantitatively important under reasonable parameter regimes (Fig. 2C). For example, if recombination is shut down along only 10% of the sex chromosome, the fitness costs that accumulate for novel sexual genotypes after 200,000 generations and under relatively weak stabilizing selection (e.g., of similar strength to that acting on human height (2)) are comparable in magnitude to the sexually antagonistic fitness benefits that have been hypothesized to drive sex chromosome turnovers (22, 23), or to the costs attributed to recessive deleterious mutations that have been proposed to underlie sex chromosome stasis (28). This suggests that the barrier presented here may be a potent source of sex chromosome stability, even when recombination shutdown affects only a portion of the sex chromosomes. Furthermore, as the non-recombining region expands over the course of sex chromosome evolution, the barrier that it presents to sex chromosome turnover accelerates in proportion to the increasing genetic variance contributed by the region (Fig. 2B), although this process might be checked by gene degeneration, and therefore reduced contribution to the trait’s genetic variance, in the nonrecombining region of the sex-specific chromosome (as discussed below).

Chromosome-specific drift is a cumulative process: the divergence in different chromosomes’ genetic values—and therefore the fitness costs to novel sexual genotypes—increases over time. The barrier to sex chromosome turnover generated by chromosome-specific drift can thus facilitate further sex chromosome differentiation: when a shutdown of recombination along a portion of the sex chromosomes hinders their turnover, this stability allows time for the accumulation of further recombination shutdowns. This mechanism does not, in isolation, generate selection for additional recombination suppression, but by stabilizing the existing sex chromosome system, it may indirectly promote extensions of the non-recombining region and facilitate the accumulation of other barriers to sex chromosome turnover. Given the ubiquity of polygenic traits, chromosome-specific drift may thus act as a global mechanism promoting sex chromosome stability across taxa, particularly in the early stages of sex chromosome evolution.

All else equal, regions of the sex chromosomes that underwent recombination shutdown earlier—that is, older ‘strata’—will have come to generate a stronger barrier to sex chromosome turnover, because they have had more time to drift apart in their genetic values and are thus associated with more extreme displacements from trait optima when rearranged in novel sexual genotypes (Fig. S3). However, these older strata are also more vulnerable to other evolutionary processes, such as gene degradation on the sex-specific chromosome (18, 29), which reduce the degree to which they contribute to phenotypes and thus decrease their importance in generating fitness costs for novel sexual genotypes. The contribution of each stratum to the barrier to sex chromosome turnover is therefore a balance between the timing of recombination shutdown and the genetic contribution of the non-recombining region to the trait.

The strength of the barrier presented by novel sexual genotypes’ fitness costs to the sex chromosome turnover can be summarized in a single ‘effective selection coefficient’, *s*_eff_, which is especially useful in cases where turnovers produce multiple novel sexual genotypes. If there are *n* + 1 total genotypes in the turnover, *s*_eff_ is the specific eigenvalue, minus 1, of the *n* × *n* matrix describing the deterministic dynamical system that corresponds to the direction, in n dimensional space, of invasion of the new sex-determining system (see Appendix Section S2; 30, 31). Monte Carlo simulation reveals that the probability of a sex chromosome turnover is well approximated by the standard fixation probability function for a single genetic variant (32), evaluated at the selection coefficient *s*_eff_ (Fig. S15), justifying the use of this metric for the barrier to sex chromosome turnover. For various kinds of sex chromosome turnover, across replicate instances of chromosome-specific drift under empirically reasonable parameter configurations, the average value of *s*_eff_ declines from an initial value of 0 to values that would drastically reduce the probability of a sex chromosome turnover within ∼100,000 generations (Fig. 2).

Because chromosome-specific drift is an inherently stochastic process, though its average effect is to cause novel sexual genotypes to be poorly adapted to the trait optimum (Fig. 2), some realizations of the process have this effect and some do not (Fig. S1). For this reason, and because the probability of fixation is a convex function for negative selection coefficients, the average probability of invasion across realizations of the fitnesses of novel sexual genotypes does not equal—and in general is higher than—the probability of invasion conditional on the average fitness of the novel sexual genotypes (or the average value of *s*_eff_) across realizations. Nonetheless, explicit simulations of the invasion process reveal that the average probability of invasion declines rapidly on realistic timescales, even in small populations (Fig. 2F–G, S4, S5).

As noted above, the rate of chromosome-specific drift under stabilizing selection is relatively insensitive to the strength of selection (27). In contrast, the fitness cost to a novel sexual genotype that, because of chromosome-specific drift, is displaced a certain distance from the trait optimum is larger if selection is stronger. Together, these two facts imply that the barrier to sex-chromosome turnover induced by chromosome-specific drift will be larger if stabilizing selection is stronger. Nonetheless, even under relatively weak stabilizing selection, the barrier generated by chromosome-specific drift can be substantial (Fig. S6), an important feature given the apparent ubiquity of weak stabilizing selection across taxa (2, 4, 33).

Additionally, chromosome-specific drift is more rapid, and will therefore be more likely to inhibit sex chromosome turnover, in smaller populations (Fig. S4, S5; Appendix Section S1). However, the consequences of genetic drift for sex chromosome evolution are complex. Drift could promote sex-chromosome turnover by enabling the transient spread of weakly deleterious sex-determining mutations, and it can even confer on these mutations an indirect selective advantage (31, 34). Given these countervailing forces, it is difficult to definitively assess the impact of population size (or subdivision) on the likelihood of sex chromosome turnover. Nevertheless, the theory described here highlights an additional pathway by which genetic drift may impede, rather than facilitate, sex chromosome turnover.

A particularly salient mechanism by which the effective strength of selection—and therefore the barriers to sex chromosome turnover—can increase is pleiotropy. Pleiotropy, where genes affect more than one phenotype, is common for highly polygenic traits (4, 35). It does not prevent chromosome-specific drift (27), and displaces novel sexual genotypes across a multi-dimensional fitness landscape, intensifying their fitness costs. This is true even under conservative assumptions (e.g., only two traits under weak stabilizing selection; Fig. S7), suggesting that even when selection on individual traits is weak, the compounding effects of selection on multiple traits with pleiotropic genetics can substantially magnify the barriers to sex chromosome turnover. More generally, these dynamics are not confined to strict pleiotropy: whenever multiple polygenic traits map to the same chromosomes, rearrangements of those chromosomes impose multi-dimensional fitness costs.

The genetic architecture of traits under selection plays a crucial role in the stability of the existing sex chromosome system in this theory. The divergence in the genetic values of the sex chromosomes (or the non-recombining region) is faster under more sex-specific genetic architectures with a weaker male–female correlation in allelic effects (Fig. S8), suggesting that sex-based specialization in the genetic architecture of polygenic traits will intensify the barrier to sex chromosome turnover. Additionally, the rate of divergence of the sex chromosomes’ values is proportional to the genetic variance contributed by their non-recombining region, which in turn is affected by the distribution of trait-affecting loci on the original sex chromosomes. Greater ‘polygenicity’—the result of higher mutation rates or larger mutational targets— facilitates chromosome-specific drift and thus enhances barriers to turnover (Fig. S5, S9, S10). However, even when the polygenicity or sex-specificity of the genetic architecture are reduced, the mechanism that I have described can generate significant barriers to sex chromosome turnover under the combined impact of selection on multiple traits (Fig. S7). Furthermore, although the results presented here assume an additive genetic basis for traits under selection (consistent with evidence on the genetic architecture of many polygenic traits; 4, 36), departures from additivity across loci (epistasis) or within loci (dominance) could further intensify the fitness costs to novel sexual genotypes and thereby reinforce the barrier to sex chromosome turnover.

Divergence in the genetic values of the sex chromosomes may also be slowed by occasional recombination in their otherwise non-recombining regions. One mechanism for such recombination is environmentally-induced sex reversal, which is common in amphibians and fish (37–39). In sex-reversed individuals, recombination patterns typically follow phenotypic rather than genotypic sex (40, 41), allowing the X and Y (or Z and W) to recombine. This will tend to equilibrate their genetic values, slowing the accumulation of barriers to sex chromosome turnover (Fig. S11). Environmental sex reversal has been proposed to explain the relatively high frequency of sex chromosome turnover in true frogs (37, 40, 41). The theory described here offers a complementary argument for how lability in sexual development, rather than being merely a corollary of developmental noise, can play a fundamental role in shaping the evolution of genetic systems.

Turnovers in the system of heterogamety that involve new dominant major sex-determining genes produce, in their course, females that carry former Y chromosomes or males that carry former W chromosomes. Such turnovers will be especially disfavored under stabilizing selection since, for example, the Y chromosome from an initial male-heterogametic system was previously carried only by males, allowing its female genetic value for the trait to have drifted unconstrainedly in the non-recombining region of that chromosome. Females who subsequently come to carry the Y will therefore deviate especially far from the trait optimum (Fig. 2, Fig. S1). The theory thus predicts that turnovers involving dominant major sex-determining genes should tend to preserve, rather than change, the system of heterogamety, a prediction that is borne out in phylogenetically rigorous analyses of sex chromosome turnovers in true frogs (42).

Because any sex chromosome turnover involves the generation of novel sexual genotypes (21), the theory presented here can explain the stability of sex chromosome systems against kinds of transitions that are permitted by other theories. For instance, turnovers in the linkage group that determines sex without a change in the system of heterogamety can occur readily under alternative theories such as those which rely on sexually antagonistic selection, but, because these turnovers produce males homozygous for the former X chromosome (or females homozygous for the former Z), they are impeded by stabilizing selection (Fig. 2). Additionally, certain heterologous transitions between male and female heterogamety do not produce YY males or WW females (12); barriers to these transitions are not predicted by theories based on large fitness costs to YY or WW individuals, but are predicted by the present theory (Fig. S12).

The mechanism proposed here will naturally act in conjunction with alternative mechanisms promoting stability of sex chromosome systems over time, such as the accumulation of deleterious mutations on the sex-specific chromosome (21, 43, 44); the presence of sexually antagonistic mutations on either sex chromosome (15, 22, 23); or the evolution of regulatory divergence between the sex chromosomes (18). Given the ubiquity of polygenic traits across taxa, however, the mechanism proposed here is likely to exert a substantial, and previously overlooked, influence on the dynamics of sex chromosome evolution, particularly during the initial stages of sex chromosome divergence when alternative mechanisms have yet to arise. Over evolutionary timescales, these early-acting effects may contribute substantially to shaping the distribution of sex-determining systems across taxa.

It is also important to note that the theory described here does not preclude sex chromosome turnovers outright. It does, however, imply that the forces driving them, such as sexually antagonistic (22, 23) or sex-differential (45) selection, must be sufficiently strong to overcome the fitness disadvantages to novel sexual genotypes under stabilizing selection. In this way, chromosome-specific drift acts as a ‘ticking clock’: once a new sex chromosome system has evolved, the sex chromosomes drift increasingly over time in their average genetic values for traits under stabilizing selection, gradually decreasing the chance that another transition will be possible. The theory thus predicts that species with young sex chromosome systems are more likely to experience another turnover, relative to species with more established sex chromosome systems. This time constraint will be particularly relevant to heterologous transitions in the heterogametic system, in which former autosomes come to act as neo-sex chromosomes, presenting a clean slate for subsequent divergence of the neo-sex chromosomes’ genetic values (Fig. 2). This temporal prediction is supported by phylogenetic evidence in vertebrates, with clades experiencing either frequent transitions or near-complete stasis (11, 46–48).

In addition to explaining barriers to turnover between sex chromosome systems, the theory can also explain the empirical preponderance of transitions from environmental sex determination (ESD) to genetic sex determination (GSD) versus the reverse (11, 49, 50). Under ESD, all chromosomes are autosomes. The transformation of one pair to sex chromosomes via a new sex-determining mutation (or mutations) does not alter the chromosomal complement of either males or females, and is therefore not disfavored by stabilizing selection. However, the reverse transition from GSD to ESD produces males homozygous for the former X chromosome or females homozygous for the former Z, and is therefore hindered by stabilizing selection.

Thus far, I have described the theory in terms of stabilizing selection around a constant optimal trait value. However, adaptive landscapes are often dynamic (51–53). If the optimal trait value shifts, the population experiences directional selection towards the new optimum, which is attained via concerted small changes in allele frequencies at many loci throughout the genome; once the new optimum is reached, stabilizing selection resumes (24, 54, 55). Thus, the average genetic values of the chromosomes all shift coordinately during the phase of directional selection (Fig. 3, S13), after which they resume the constrained drift process described above. Therefore, the fitness disadvantages suffered by novel sexual genotypes are not, in the long run, impacted by the shift in trait optimum. However, these disadvantages are transiently reduced during the period of directional selection (Fig. 3), suggesting that sex chromosome turnovers may be more likely immediately following environmental disturbances or other events that reshape the fitness landscape. Environmental or ecological changes that elicit adaptive responses in quantitative traits can therefore indirectly promote transitions in genetic sex-determining systems in this model—an intriguing potential side effect of the sustained environmental shifts expected under climate change.

**Fig. 3.**
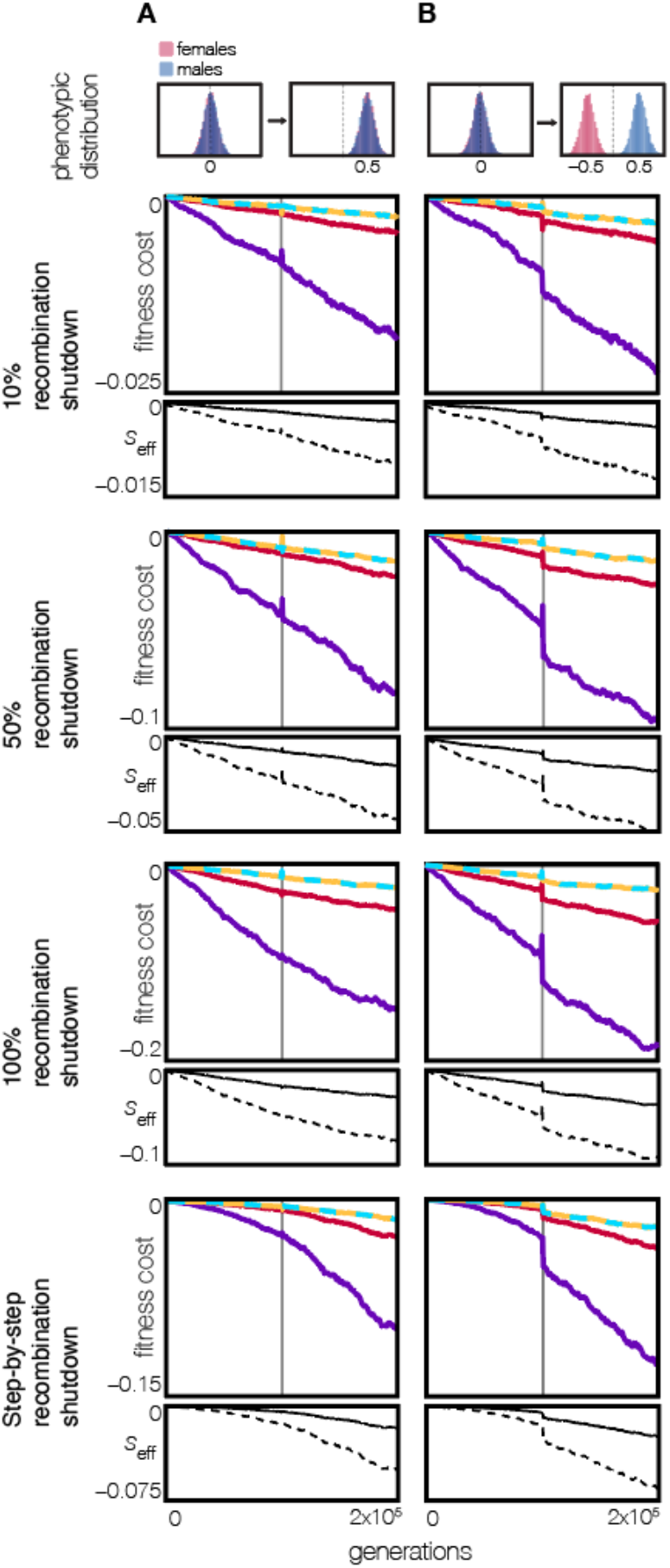
Sexually antagonistic shifts in fitness optima exacerbate barriers to sex chromosome turnover, while sexually concordant shifts do not. **(A)** When a trait’s optimum shifts in the same direction for males and females, the genetic values of all chromosomes shift in concert in the same direction (Fig. S13), and so there is no lasting effect of the shift on the fitness of novel sexual genotypes and therefore the strength of the barrier to sex chromosome turnover, relative to the case of no optimum shift. **(B)** If the trait optimum shifts in different directions for males and females, the X and Y chromosomes uniquely and disproportionately contribute to evolution towards these new optima (26) (Fig. S14). This causes a lasting reduction in the fitness of novel sexual genotypes, which involve imbalances of the original X and/or Y, and a concomitant strengthening of the barrier to sex chromosome turnover. In both sexually concordant and antagonistic optimum shifts, however, the fitness costs to novel sexual genotypes can be transiently lowered in the generations immediately following the optimum shift, suggesting that these brief periods may be especially conducive to sex chromosome turnover. Results shown are from simulations in which the extent of recombination suppression varies from 10%, 50%, and 100% recombination shutdown between the X and Y chromosomes; or a case in which recombination is suppressed in successive increments of 10% of the length of the chromosome every 2*N* generations. Note that, to facilitate comparison between the sexually antagonistic and concordant shifts in fitness optima, the vertical scale varies based on the recombination scenario.

Of particular interest is the case where the optimum shifts differently for males and females, inducing sexually antagonistic selection and ultimately sexual dimorphism in the trait (Fig. 3). Sexually antagonistic optimum shifts provoke disproportionate sex-specific changes in the genetic values of the sex chromosomes relative to the autosomes. Under male heterogamety, the Y exclusively promotes evolution towards the new male optimum, while the X predominantly promotes evolution towards the new female optimum (26) (Fig. S14). The sex chromosomes (specifically their non-recombining regions) therefore come to contribute uniquely and disproportionately to the maintenance of male and female trait optima (Fig. 3, S13), exacerbating the fitness costs suffered by novel sexual genotypes during a sex chromosome transition, as chromosomes that have evolved to promote one sex’s trait optimum find themselves in the other sex. Moreover, since this effect is a result of directional selection on, rather than drift of, chromosomes’ genetic values, it is established on a much faster timescale. Sex chromosome stability has traditionally been studied as a cause of sexual dimorphism (56, 57). Notably, the effect just described suggests a reverse causality, with the outsized role of the sex chromosomes in the evolution of sexual dimorphism stabilizing the sex chromosome system. Teasing apart these directions of causality may soon be possible via phylogenetic methods applied to new, large-scale data on sex-determining mechanisms across diverse taxa (11, 46, 47, 58, 59).

The theory proposed here is similar in kind to theories of ‘developmental system drift’ (60, 61) where, for example, the individual components of a developmental system (62, 63) or regulatory network (64) can undergo evolutionary change while together maintaining a constant phenotypic effect. An advantage of the present theory is that it is cast in concrete population genetic terms— genetic variances, population sizes, selection strengths—so that its consequences, such as for the rate of accumulation of fitness disadvantages to novel sexual genotypes, can be analyzed using the machinery of modern population genetic theory. Chromosome-specific drift may therefore offer a useful framework for investigating the dynamics of polygenic traits across diverse evolutionary contexts (27).

## Acknowledgments

I thank C. Veller for research assistance and comments on the manuscript. I am grateful to I. Qureshi for comments on the manuscript and G. Coop, J. Berg, M. Steinrücken, L. Delph, M. Hahn, and A. Dahl for helpful discussions. I am also grateful to B. Haller and M. Jarsulic for advice on simulations. The simulations in this paper were run on the Randi computing cluster, supported by the Center for Research Informatics in the Biological Sciences Division at the University of Chicago, and the Midway 3 computing cluster, supported by the Research Computing Center at the University of Chicago.

## Methods

### Simulations

#### Main model

The population is composed of *N* = 10,000 diploid individuals (5,000 males and 5,000 females) and evolves according to a Wright–Fisher process with random mating. Stabilizing selection is implemented via a Gaussian fitness function with a standard deviation normalized to 1. Unless otherwise stated, males and females have the same optimal phenotype, arbitrarily coded as 0. Under the simulation parameters described below, the strength of stabilizing selection, measured as the ratio between the standard deviation of the fitness function and the standard deviation of the phenotypic distribution, is approximately 5. This strength is similar to empirical estimates of the strength of stabilizing selection on quantitative traits in humans (2, 4, 66) and other taxa (33).

An individual’s genome is composed of *L* = 1,000 loci, approximating a quantitative phenotype controlled by 1000 ‘causal’ loci. These 1,000 loci are evenly distributed across 10 chromosomes (9 autosomes and an X and Y chromosome pair). Recombination occurs freely between chromosomes (at rate *r* = 0.5). The recombination rate between adjacent loci on the autosomes in both sexes and the X chromosome in females is *r* = 0.01, so that each chromosome experiences one crossover per gamete on average. There is no crossover interference.

The genetic basis of the quantitative trait is additive, such that an individual’s trait value is the sum of the phenotypic effects of all of the alleles that they carry. Unless otherwise stated, allelic variation arises in the simulations via mutations, which occur at rate *μ* = 10^−5^ per locus per generation, resulting in a genome-wide mutation rate of *U* = *Lμ* = 0.01 per generation. The male and female phenotypic effects of new mutations are drawn from a bivariate normal distribution with mean 0, standard deviation 0.05, and correlation *ρ* = 0.9, so that the phenotypic effects of alleles in males and females are highly, but not perfectly, correlated (6–10)

Simulations were initialized with a burn-in period of 10*N* generations, during which the X and Y chromosomes recombined freely along their entire length, at a rate equivalent to the autosomal recombination rate (so that the sex chromosomes initially functioned as entirely pseudo-autosomal regions). Following the burn-in, recombination suppression was imposed according to four distinct scenarios, designed to reflect different stages or modes of sex chromosome evolution. In all cases, the sex-determining mutation was positioned at the distal end of the focal chromosome and was included within the non-recombining region.

In scenarios (a)–(c), recombination between the X and Y chromosomes was immediately and permanently suppressed along 10%, 50%, or 100% of their length, respectively. Scenario (d) modeled a gradual recombination shutdown. Starting immediately after the burn-in, recombination suppression expanded in 10% increments every 20,000 (i.e., 2*N*) generations until the entire chromosome became non-recombining. Although the tempo and mechanism of recombination suppression (e.g., inversion accumulation) vary across taxa, this scenario provides a stylized representation of progressive recombination loss across the sex chromosomes.

All calculations were performed relative to the state of the population at the end of this burn-in period. For all fitness calculations, the fitness disadvantage of individuals with novel sexual genotypes is calculated relative to the fitness of individuals with the original sexual genotypes (XY males and XX females).

Simulations were carried out in SLiM 4.3 (67). Unless otherwise stated, results shown in the Main Text and supplementary figures are averaged across 250 independent replicates. Simulation code is available at: https://github.com/Pavitra451/polygenic_barriers_to_sex_chromosome_turnover/.

#### Optimum shift

In simulations which involve a shift in fitness optimum, an instantaneous shift in the fitness optima of males and females occurs 10*N* generations after the conclusion of the burn-in period. This shift was either concordant between the sexes, with the male and female fitness optima changing to the same value of 0.5, or sexually antagonistic, with the male optimum increasing to 0.5 and the female optimum decreasing by the same amount to –0.5.

#### Environmental sex reversal

In simulations examining the effects of environmental sex reversal, 0.5% of genetic males and 0.5% of genetic females undergo sex reversal per generation, such that their genotypic sex does not correspond to their phenotypic sex as an adult. These individuals were selected randomly and suffered no fitness disadvantage independent of their value of the trait under stabilizing selection. In accord with common observation (39–41), recombination patterns in sex-reversed individuals were assumed to follow their phenotypic, not genotypic, sex. Thus, for example, the X and Y chromosomes could recombine in sex-reversed XY females.

#### Multiple traits

To model a scenario where multiple traits under selection are affected by closely linked genes on each chromosome, I implemented two distinct mutational processes across all 1,000 loci. Both mutational processes followed the parameters described above, but the first type affected only phenotype 1, while the second type affected only phenotype 2. This simulates a case in which mutations at a given gene may affect two different phenotypes. This scenario can also be generalized to a case in which mutations at different, but closely linked, genes impact different phenotypes.

### Introduction of new sex-determining mutations

Due to computational constraints, simulations testing the invasion of new sex-determining alleles were conducted with a reduced population size of *N* = 1,000. To maintain a rate of chromosome-specific drift comparable to that in the full-sized simulations, the genome-wide mutation rate per site was reduced to u = 10^−6^. This adjustment preserves the ratio of equilibrium genetic variance to the population size, and thus preserves the rate of chromosome-specific drift between the two simulation setups (Fig. S5; Appendix Section S1).

To calculate the effects of chromosome-specific drift on invasion probabilities, simulations were conducted in two sequential phases. The first phase replicated the simulation scenarios described above, but with the smaller population size. A burn-in period of 100,000 generations (100*N*) was used, during which no recombination shutdown occurred. Following this burn-in, recombination was shut down along either 10% or 100% of the sex-specific chromosome, in a region including the sex-determining mutation. The simulation “state”—which contains all information necessary to fully restart the simulation—was saved at multiple timepoints: 0, 10, 100, 1,000, 10,000, 100,000, 200,000, 300,000, 400,000, and 500,000 generations after the end of the burn-in. For each parameter combination, 500 replicate simulations were run and saved.

In the second phase, for each saved timepoint, the simulation was reloaded from its saved state, and a new dominant sex-determining mutation was introduced—either on a different chromosome pair (representing a heterologous transition) or on the original sex chromosome pair (representing a homologous transition). Both male- and female-determining mutations were considered. This set of new sex-determining mutations thus encompasses sex-determiners that could facilitate all four sex chromosome turnovers shown in Fig. 2.

### Each simulation was then run until one of three outcomes occurred

(1) the new sex-determining mutation was lost from the population; (2) a sex chromosome turnover occurred— defined by the new sex-determining mutation reaching 25% frequency and triggering the fixation or loss of the ancestral sex-determining mutation; or (3) 20*N* generations elapsed without either outcome occurring. Outcome (3) was extremely rare, occurring only 1 time across 800,000,000 total simulations.

Using 500 saved starting configurations and 20,000 replicate simulations per starting configuration, this approach generates total of 10 million replicate introductions of a sex-determining mutation for each of the 80 parameter combinations (2 recombination shutdown regimes; 4 types of possible transitions; 10 time points) shown in Fig. 2F-G.

### Calculating the effective selection coefficient against turnover, and estimating the turnover probabilities, for various configurations of the fitnesses of novel sexual genotypes

Here, I describe the scheme used to analyze the dependence of the probability of sex-chromosome turnover on the fitnesses of the novel sexual genotypes, and to show that this dependence is well captured by a certain eigenvalue of the dynamical system.

For both homologous and heterologous transitions, I generated a large number of configurations of the fitnesses of the novel sexual genotypes produced in the course of each type of turnover. To avoid scenarios where some novel sexual genotypes are beneficial while others are deleterious, which could lead to intermediate equilibria and therefore hang simulations, in each configuration it was decided randomly, with even probability, whether the selection coefficients on the novel sexual genotypes would be all negative or all positive. For 500 configurations, a maximum selection coefficient *s*_max_ was drawn from a uniform distribution on [-0.01, 0] (for negative selection coefficients) or on [0, 0.01] (for positive selection coefficients); for 500 configurations it was drawn from a normal distribution with mean 0 and standard deviation 0.005, discarding values outside of [–0.01, 0.01]. This “mixed” sampling scheme was used to ensure a reasonable coverage of the interval [–0.01, 0.01] for the effective selection coefficient *s*_eff_ described below. Having drawn *s*_max_ for a particular configuration, selection coefficients were drawn independently for each novel sexual genotype from a uniform distribution on [*s*_max_, 0] if *s*_max_ < 0 or on [0, *s*_max_] if *s*_max_ > 0.

Having chosen the selection coefficients on the novel sexual genotypes, the eigenvalues and eigenvectors of the resulting Jacobian matrix of the dynamical system were calculated (see Appendix). The eigenvalue λ_inv_ was identified that corresponded to the eigenvector closest (in Euclidean distance) to the eigenvector of the neutral system that points in the direction of the neutral invasion path: these eigenvectors are [0.8729, –0.4364, 0.2182] for homologous transitions and [0.8528, –0.2132, –0.4264, 0.2132] for heterologous transitions, and both correspond to an eigenvalue of 1 in their respective neutral systems (27). The effective selection coefficient for the configuration was taken to be *s*_eff_ = λ_inv_ – 1.

Then, for each configuration of the selection coefficients on the novel sexual genotypes, I simulated 10^7^ Wright–Fisher trials of a randomly mating population of size 1,000. Simulations were carried out in MATLAB R2023b. Each trial began with a single copy of the new dominant female-determining mutation X′ (for homologous transitions) or A′ (for heterologous transitions). For homologous transitions, this mutation first appeared in an X′X or an X′Y female, with equal probability; for heterologous transitions, the mutation first appeared in an XX A′A or an XY A′A female, again with equal probability. In both cases, each trial began with exactly 500 females and 500 males. From these starting points, each trial was iterated until only the two sexual genotypes associated with the original male-heterogametic system, or of the new female-heterogametic system, remained; trials in which the latter occurred were scored as “fixations”. The transition probability for the configuration was then estimated as the proportion of fixations out of the 10^7^ trials.

Fig. S15 plots *s*_eff_ calculated for the various sampled configurations of fitnesses of novel sexual genotypes against the fixation probabilities estimated for these various configurations. It can be seen that the fixation probability is, for both homologous and heterologous transitions, a near perfectly monotonic function of *s*_eff_, which in fact closely corresponds to the standard formula (32) for the fixation probability of an allele of selection coefficient *s*, (1 – *e*^−2*s*^)/(1 – *e*^−4*Ns*^).

### Analysis of human height

The chromosome-specific additive genetic values displayed in Fig. 1b were calculated using summary statistics from the meta-analysis of human height GWAS by Yengo et al. (65), downloaded from the Giant Consortium website (https://portals.broadinstitute.org/collaboration/giant/index.php/GIANT_consortium_data_files; data file labelled ‘Yengo.2022.height.GWAS.ALL’). The X chromosome was excluded from these analyses. The estimated effect of each variant was averaged across males and females.

The polygenic score (PGS) of each autosome was calculated as the sum of the effects of the variants contained on the autosome, weighted by their frequencies. The PGS for each autosome was then normalized by the genome-wide value, to identify relatively height-increasing and height-decreasing chromosomes. Note that, because this analysis is based on GWAS effect sizes, it takes into account only the effects of variants that are segregating in the sampled populations; it does not include fixed variants, which also contribute to differences across chromosomes in their mean genetic values, or variants not segregating in the sampled populations but segregating in other populations. The differences between chromosomes displayed in Fig. 1b is therefore likely an underestimate of the true differences across human autosomes in their genetic values for height.

## Notes

### Competing Interest Statement

The authors have declared no competing interest.

### Summary of Updates

Analyses extended and updated; Figs. 1-3 and main text revised.

