## Supplementary Figures and Appendix for "Chromosome-specific drift under stabilizing selection generates polygenic barriers to sex chromosome turnover"

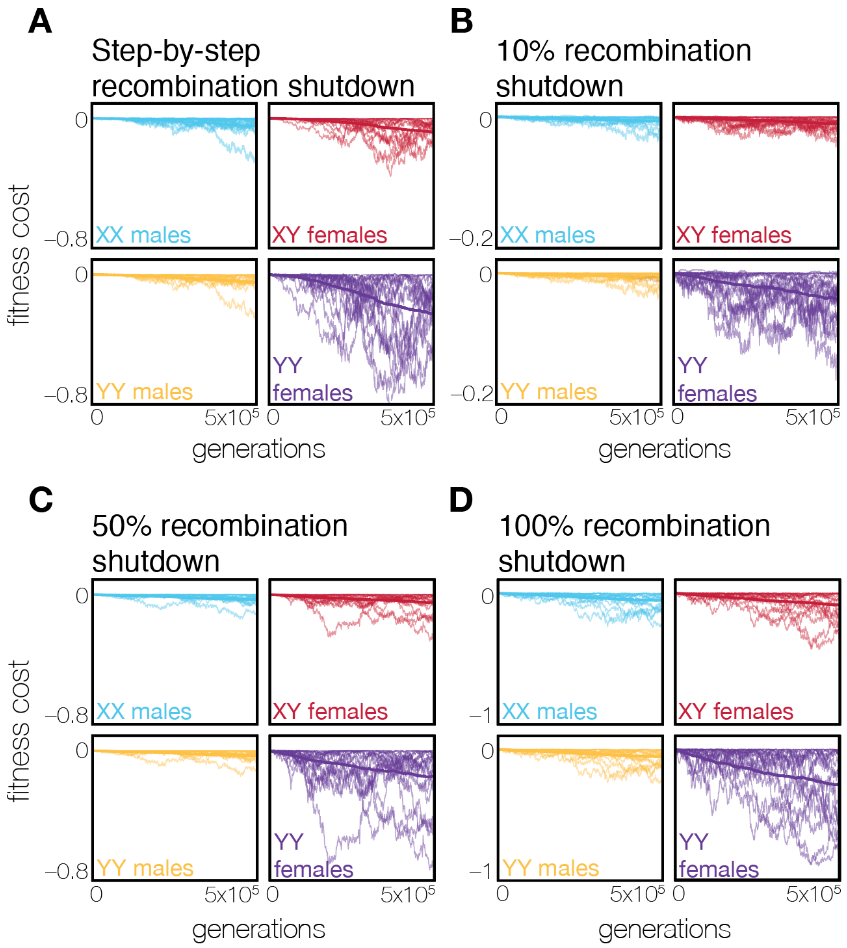


**Fig. S1. The fitness costs associated with novel sexual genotypes reflect the stochasticity of chromosome-specific drift.** Results are shown for simulations in which recombination shuts down gradually across the X and Y chromosomes (A), or in which 10% (B), 50% (C), or 100% (D) of the X and Y chromosomes are non-recombining throughout the simulation (see Methods for details). Each faint line represents the relative fitness reduction to novel sexual genotypes in a single simulation run; 20 independent simulation runs are shown per panel. Bold lines indicate average fitnesses of the sexual genotypes across 250 simulation runs. While the fitness costs to novel sexual genotypes in a particular simulation run are noisy, reflecting the drift-based divergence in each chromosome's genetic value, the average effect of this chromosome-specific drift is to decrease the fitness of novel sexual genotypes.

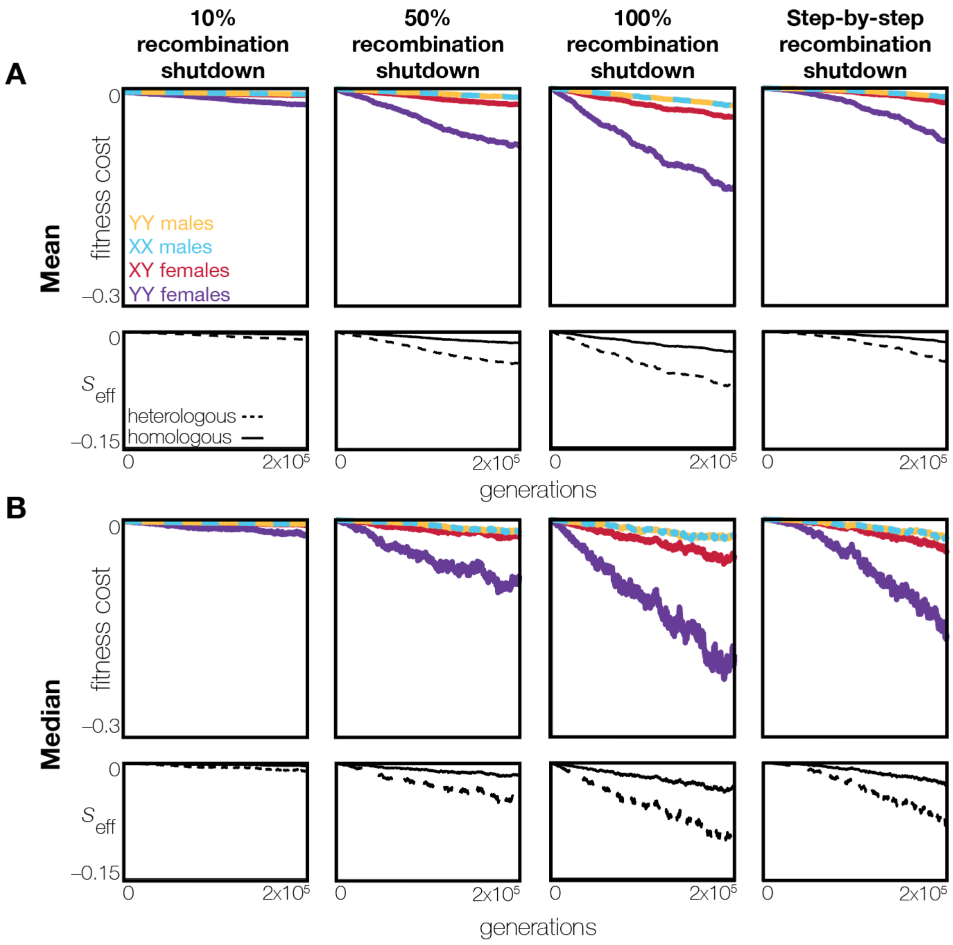


**Fig. S2. Measurement of the average barrier to sex chromosome turnover is not sensitive to the central summary statistic used (mean or median).** Panel (A) corresponds to the scenario displayed in Fig. 2B–D of the main text, in which the mean fitness cost of new sexual genotypes is calculated by averaging across 250 replicates. Panel (B) uses the same simulations but displays the median fitness cost of new sexual genotypes across 250 replicates.

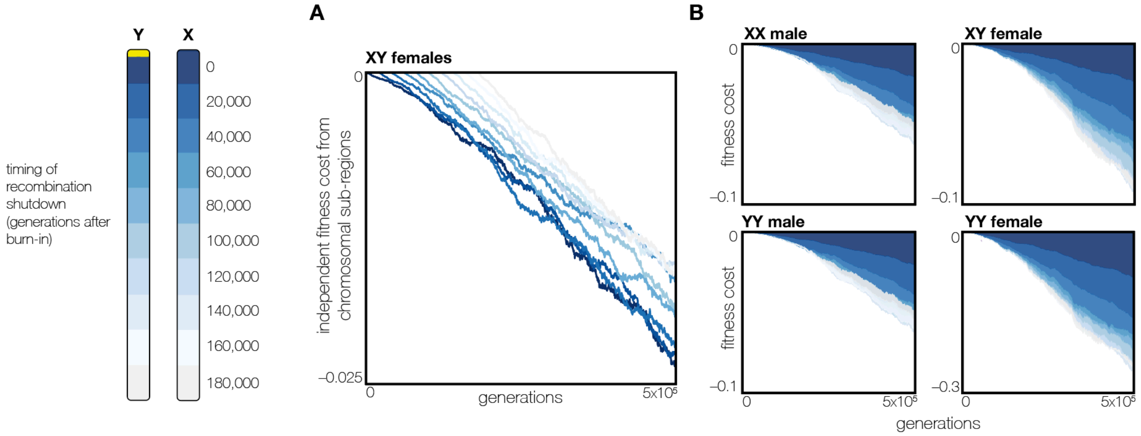


**Fig. S3. Sex chromosome strata diverge in their genetic values at similar rates following recombination suppression.** Parameters are as in Fig. 2B, with the sex chromosome undergoing gradual recombination suppression over 2N generations ($N$ = 10,000). The yellow bar indicates the position of the male sex-determining mutation. (A) As recombination shuts down sequentially across different regions of the sex chromosome, the X and Y gene homologs within each region (or 'stratum') accumulate differences in genetic values due to drift. This divergence occurs at a comparable rate in all strata, regardless of the timing of recombination suppression. Here, the independent contributions of different strata to the fitness costs of XY females is shown. (B) Although the rate of divergence between X and Y regions is consistent across different strata, those strata that experienced recombination shutdown earlier— 'older' strata— have accumulated greater genetic differences between their X and Y homologs. As a result, these older regions contribute more substantially to the overall fitness costs associated with novel sexual genotypes compared to 'younger' strata, where recombination suppression happened more recently.

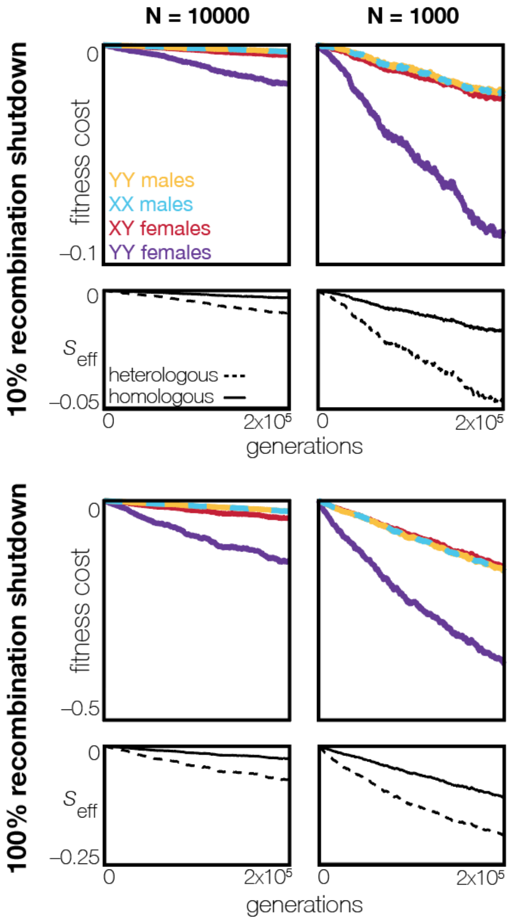


**Fig. S4. Smaller population sizes accelerate chromosome-specific drift, more rapidly impeding sex chromosome turnovers.** Parameters are as in Fig. 2 in (A), but the population size is reduced to $N = 1{,}000$ in (B).

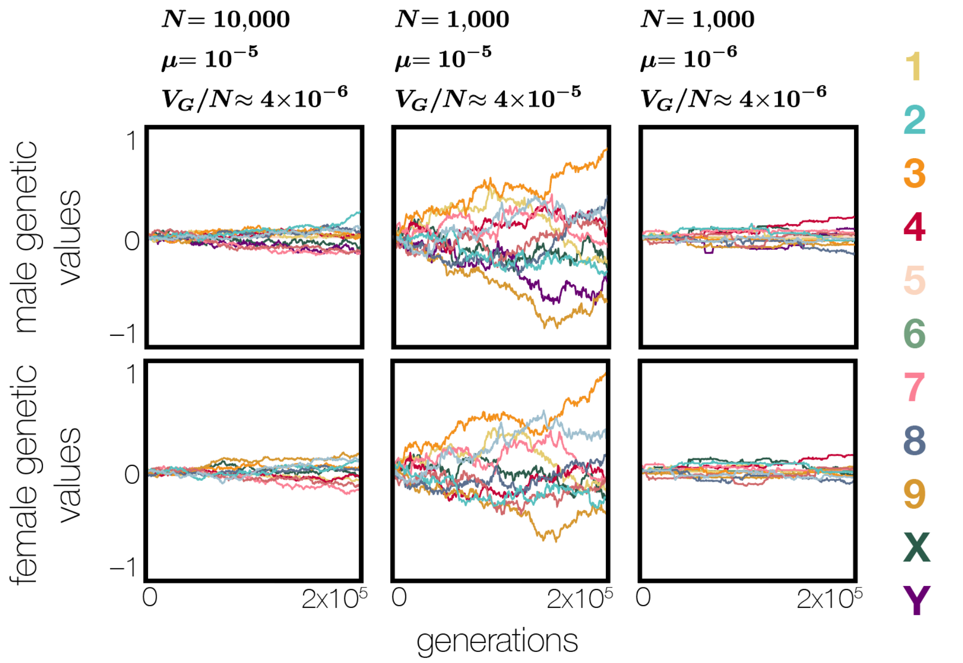


**Fig. S5. Chromosome-specific drift occurs at similar rates in populations with the same ratio of genetic variation to effective population size.** The rate of chromosome-specific drift is proportional to $V_G/N_e$, where $V_G$ is the genetic variance for the trait and $N_e$ is the effective population size (see Appendix Section S1). Under stabilizing selection, at mutation–selection–drift balance in the large-population limit, $V_G \approx 4L\mu V_S$, where $V_S$ is the width of the selection function (= 1 in my simulations), $L$ is the number of loci at which mutations affect the trait's value (= 1,000 in my simulations), and $\mu$ is the per-locus mutation rate. In panel (A), parameters are as in Fig. 1, and, in particular, $N = 1{,}000$ and $\mu = 10^{-5}$. In panel (B), the population size is reduced to $N = 1{,}000$ while the mutation rate is kept at the value $\mu = 10^{-5}$. The smaller population size, all else equal, accelerates chromosome-specific drift since the equilibrium genetic variance $V_G \approx 4L\mu V_S$ is unchanged (since $L\mu$ is unchanged relative to panel (A)). In panel (C), the population size is $N = 1{,}000$ as in panel (B), but the mutation rate is also reduced tenfold to $\mu = 10^{-6}$, so that both the equilibrium genetic variance $V_G \approx 4L\mu V_S$ and the population size $N$ are reduced tenfold relative to panel (A). Correspondingly, the rate of chromosome-specific drift observed in panels (A) and (C) is comparable, despite the lower population size in panel (C). This correspondence allows the invasion analyses in the Main Text (Fig. 2E–F), which is computationally costly, to use lower population sizes with mutation rates adjusted to compensate; these analyses would be computationally infeasible with a larger population size of $N = 10{,}000$.

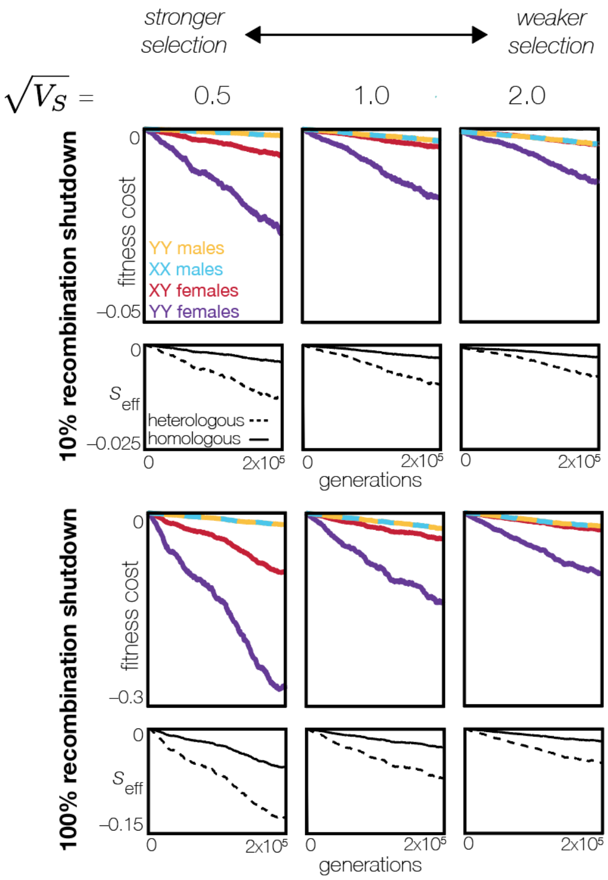


**Fig. S6. Stronger stabilizing selection on quantitative traits more rapidly impedes sex chromosome turnovers**. Parameters are as in Fig. 2, but here, results are shown for different strengths of selection, parameterized by the width $\sqrt{V_s}$ of the Gaussian fitness function (in Fig. 2, $\sqrt{V_s} = 1$). Lower values of $\sqrt{V_s}$ correspond to stronger stabilizing selection, which is seen to increase the fitness costs associated with novel sexual genotypes and therefore to decrease the probability of sex chromosome turnover. However, even weak stabilizing selection (e.g., $\sqrt{V_s}$ = 2) can generate significant barriers to sex chromosome turnover on the same time scale.

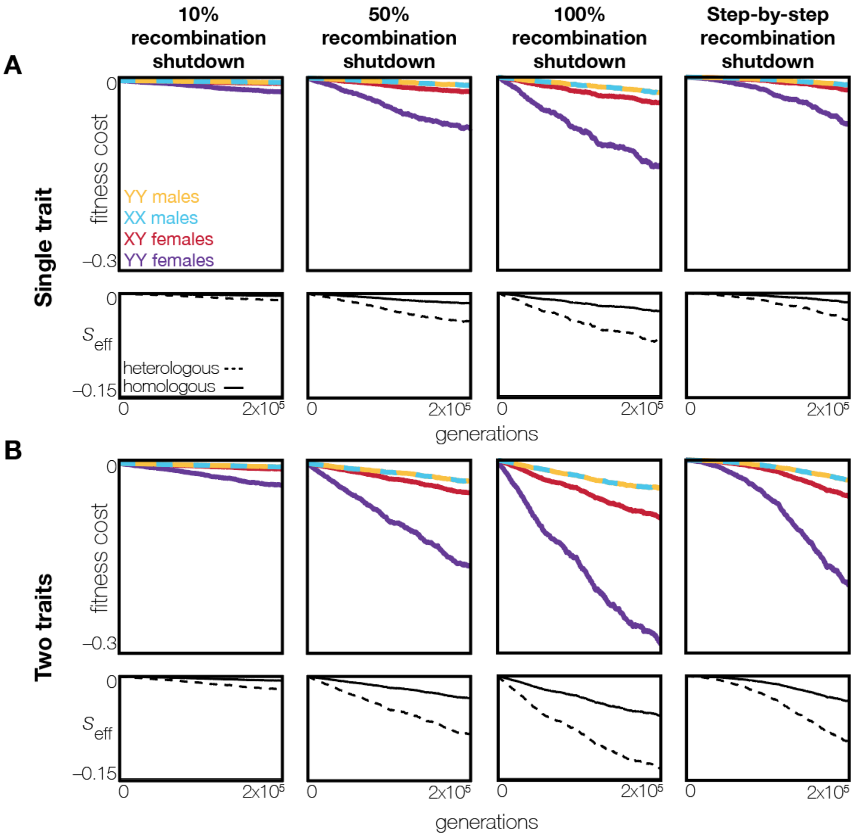


**Fig. S7. Stabilizing selection on multiple traits enhances the barrier to sex chromosome turnover.** Panel (A) corresponds to the scenario displayed in Fig. 2B–D in the Main Text, in which a single polygenic trait is encoded by 1,000 loci and experiences stabilizing selection under a Gaussian fitness function of width $\sqrt{V_s} = 1$. Panel (B) shows the fitness reductions to novel sexual genotypes in the scenario where two polygenic traits experience the same strength of stabilizing selection $\sqrt{V_s} = 1$) and are independently encoded by the same 1,000 loci (or, equivalently, by 1,000 pairs of tightly linked loci). The consequences for sex chromosome turnover of having two traits, rather than one, under stabilizing selection are approximately the same as doubling the strength of selection on a single trait. Given the ubiquity of stabilizing selection on polygenic traits, the compounding selective disadvantages arising from multiple traits under selection are likely to be a significant factor promoting barriers to sex chromosome turnover.

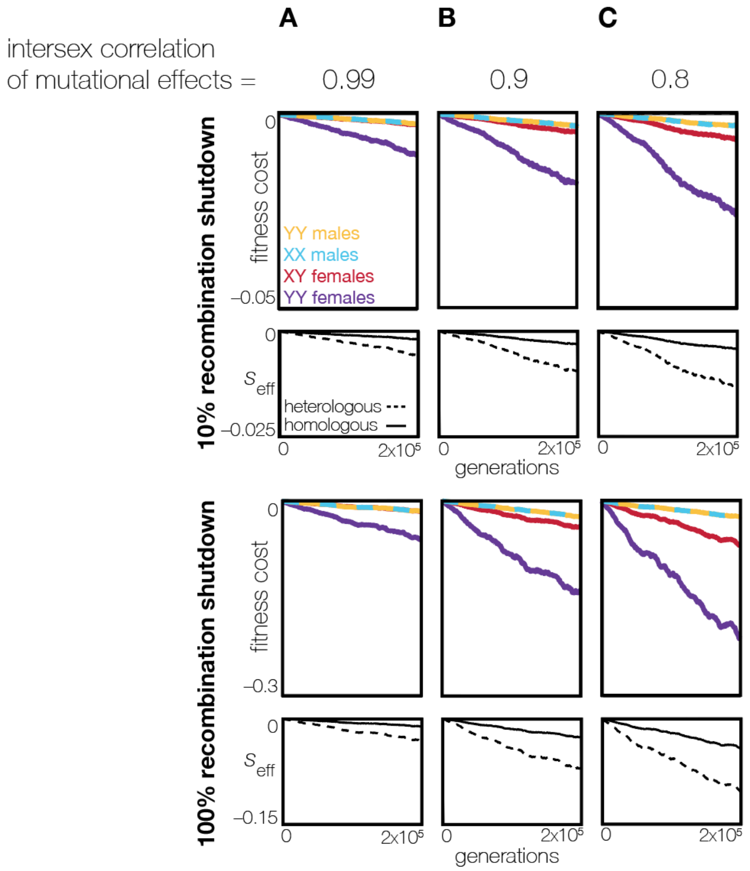


**Fig. S8. Higher sex-specificity in the genetic architecture of quantitative phenotypes enhances barriers to sex chromosome turnover**. Parameters are the same as in Fig. 2, but the correlation between the phenotypic effects of new mutations in males and females is (A) 0.99, (B) 0.9 (as in Fig. 2), or (C) 0.8, reflecting an increasing degree of sex-specificity in the genetic basis of the trait. A strongly shared genetic architecture for the trait in males and females, as in (A), constrains the genetic values of the X and Y chromosomes to resemble each other, reducing the impact of chromosomal re-arrangements and decreasing the selective disadvantages of novel sexual genotypes. In contrast, a more sex-specific genetic architecture, as in (C), allows for greater divergence between the X and Y, generating stronger barriers to sex chromosome turnover.

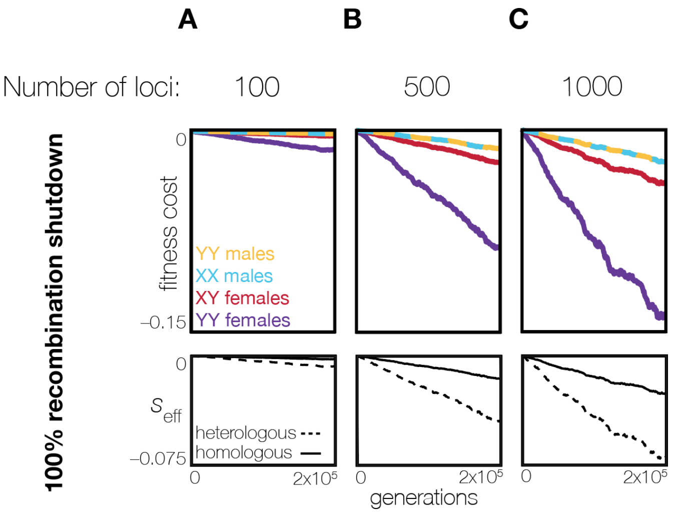


**Fig. S9. Increased polygenicity of a quantitative trait intensifies the genetic barrier to sex chromosome turnover.** Parameters are as in Fig. 2, but the number of loci contributing to the trait is (A) 100, (B) 500, or (C) 1000 (identical to Fig. 2E). The per-locus mutation rate is $\mu = 10^{-5}$ in each case.

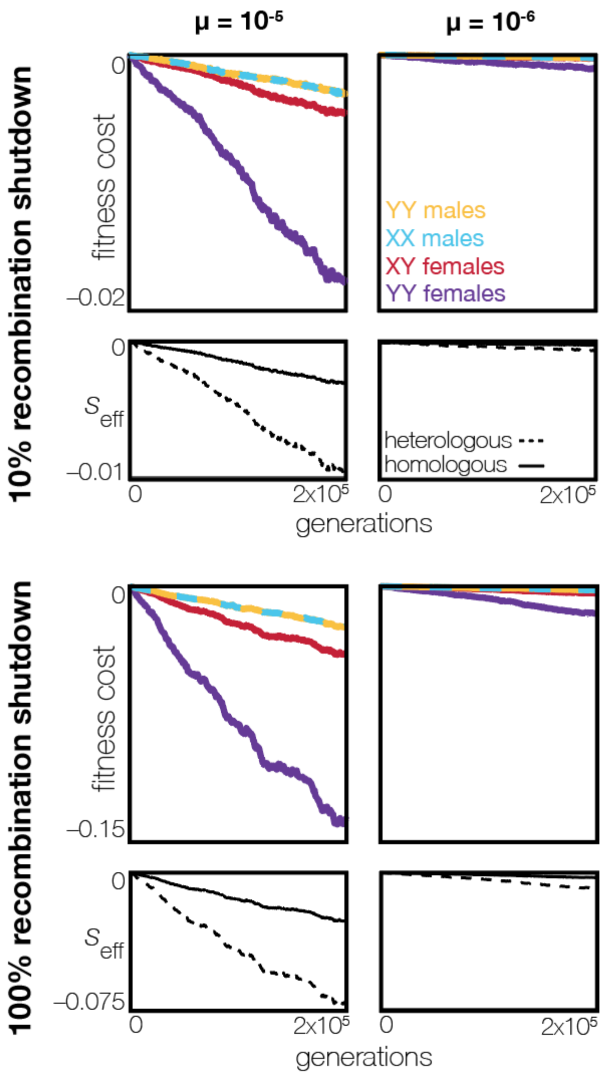


**Fig. S10. Higher mutation rates accelerate chromosome-specific drift.** Parameters are the same as in Fig. 2, except that the per-site mutation rate is reduced by an order of magnitude to $\mu = 10^{-6}$ in (B). This reduction in mutation rate lowers the equilibrium genetic variance contributed by each chromosome, thus diminishing the rate and impact of chromosome-specific drift (see Fig. S5 for the interaction between mutation rate and effective population size).

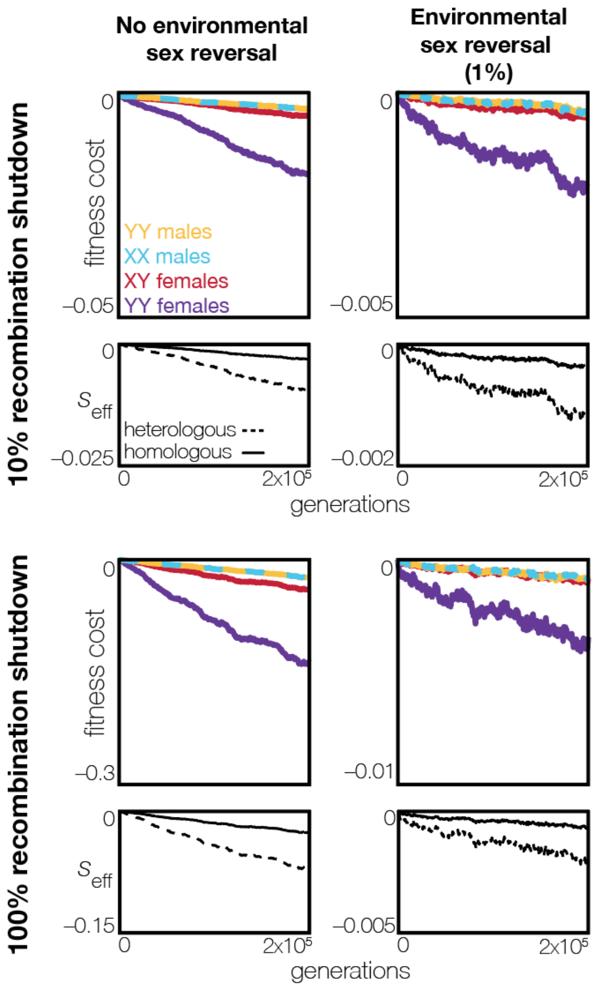


**Fig. S11. Environmental sex reversal reduces the barrier to sex chromosome turnover across extended time scales.** Parameters in panel (A) are as in Fig. 2, but in panel (B) a 1% rate of environmental sex reversal is implemented, such that 1% of adult individuals have a phenotypic sex that is discordant with their genotypic sex. This sex reversal facilitates occasional recombination between the X and Y in XY females, whose recombination pattern is based on phenotypic, rather than genotypic, sex. This occasional recombination constrains the divergence in genetic values between the sex chromosomes, which in turn slows the accumulation of fitness costs associated with novel sexual genotypes and thus prolongs the period during which a sex chromosome turnover is possible. Note that, for visual clarity, panels (A) and (B) have different y-axes.

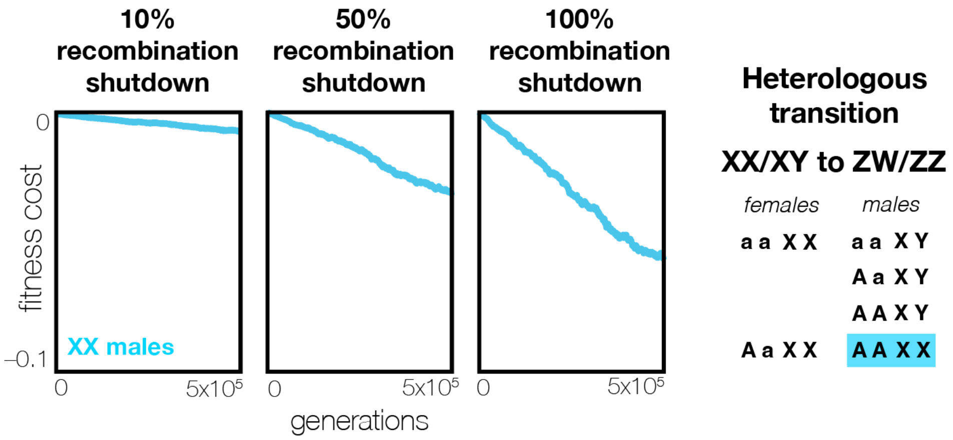


**Fig. S12. Heterogametic transitions that do not involve the creation of YY individuals are nonetheless impeded due to chromosome-specific drift.** The scenario is as in Fig. 2, but here a recessive heterologous transition in heterogamety is shown (rather than a dominant heterologous transition in heterogamety, as in Fig. 2A). This turnover is an example of a transition in sex determination that does not involve the production of YY individuals, and therefore would not be impeded by the exposure of deleterious recessive mutations that have accumulated on the Y chromosome. However, recessive heterologous turnovers are still impeded under the theory presented here, because novel sexual genotypes, which disrupt inter-chromosomal co-evolution, are necessarily produced in such turnovers.

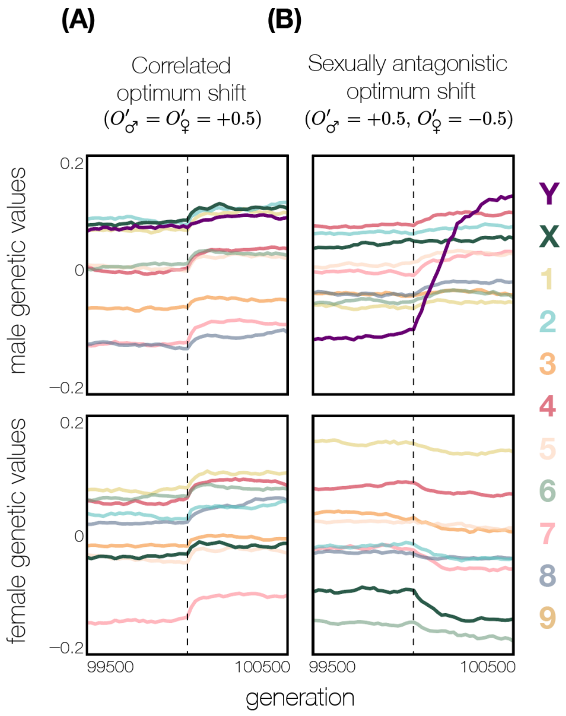


**Fig. S13. The X and Y chromosomes respond uniquely to a sexually antagonistic shift in male and female fitness optima.** Each line represents the genetic value of a chromosome in a single simulation run. (A) In response to a concordant shift in the fitness optima (here, both increased), the mean values of all chromosomes, including the X and Y, shift in concert to move the male and female phenotype to the new, increased optimal value. (B) In response to a sexually antagonistic shift in the fitness optima, the mean genetic values of the X and Y chromosomes change disproportionately in opposite directions to facilitate the evolution of, respectively, females and males to their new divergent fitness optima.

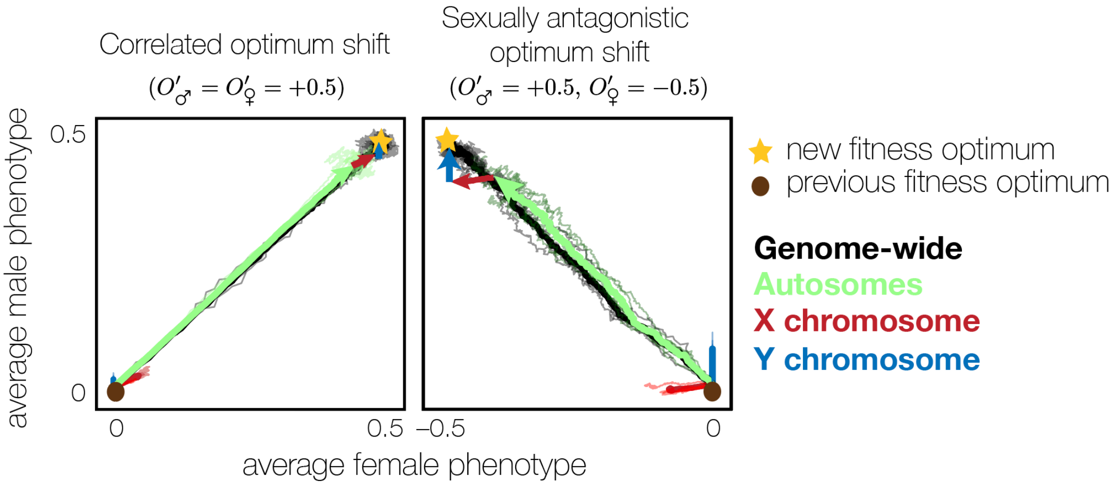


**Fig. S14. The sex chromosomes contribute disproportionately to the evolution of sexual dimorphism in quantitative traits.** The phenotypic response to selection is shown on the male–female phenotypic landscape after a concordant shift (left) and a sexually antagonistic shift (right) in the male and female fitness optima. All parameters are the same as in Fig. 3. The brown circle marks the original fitness optimum, while the yellow star marks the new fitness optimum. Each faint line represents a single replicate (5 in total) in which phenotypes were recorded every generation for 500 generations following the shift in optimum; bold lines represent the average trajectory across those 5 replicates. Trajectories are normalized relative to their values at the time of the shift in optimum, so that all trajectories start at (0,0) in male–female phenotype space. The female genetic values for the trajectory of the Y chromosome are constrained to 0 for illustration, because the Y will not normally contribute to phenotypic evolution in females in an XX/XY system. While overall phenotypic evolution (black) proceeds along a straightforward trajectory to the new optimum following both shifts, the X (red) and Y (blue) chromosomes show nearly sex-specific contributions to phenotypic evolution after the sexually antagonistic shift, in striking contrast to their concordant behavior after the concordant shift. This disproportionate contribution of the sex chromosomes to phenotypic evolution under sexually antagonistic selection drives the increase in fitness costs associated with novel sexual genotypes after a sexually antagonistic shift, as shown in Fig. 3.

### Appendix

#### S1 Chromosome-specific drift under stabilizing selection

The dynamics of chromosome-specific drift, both in the presence and absence of sex chromosomes, are derived in Veller and Muralidhar (2025). For clarity, we present here the equations for the general dynamics of chromosome-specific drift, to illustrate the underlying principles, and the particular dynamics in the presence of sex chromosomes. The latter dynamics do not explicitly model mutation, unlike in my simulations, and are therefore most precisely interpreted as describing short-term changes in the mean contributions of the sex chromosomes and autosomes to a trait under stabilizing selection (see detailed discussion in Veller and Muralidhar 2025).

##### S1.1 General Dynamics

In a diploid, random-mating population of size $N$, a trait is under Gaussian stabilizing selection of strength $V_S$ around an optimal value of zero. The genetic and phenotypic variance of the trait are $V_G$ and $V_P$.

Let the mean genetic value of a focal chromosome at time $t$ be $x_t$, and the mean genetic value of the rest of the genome be $y_t$, so that the overall mean genetic value is $z_t = x_t + y_t$. If the focal chromosome accounts for a fraction $f$ of the total genetic variance $V_G$, with the rest of the genome accounting for the remaining fraction $1 - f$, then $x_t$ and $y_t$ evolve according to the coupled stochastic differential equations:

$$dx_t = -\frac{fV_G}{V_S + V_P} z_t dt + \sqrt{\frac{fV_G}{N}}\, dW_t^1, \tag{S.1}$$

$$dy_t = -\frac{(1-f)V_G}{V_S + V_P} z_t dt + \sqrt{\frac{(1-f)V_G}{N}}\, dW_t^2, \tag{S.2}$$

where $W_t^1$ and $W_t^2$ are independent standard Brownian motions.

Notice that, from Eqs. (S.1) and (S.2),

$$\begin{aligned} dz_t = dx_t + dy_t &= -\frac{V_G}{V_S + V_P} z_t + \sqrt{\frac{fV_G}{N}}\, dW_t^1 + \sqrt{\frac{(1-f)V_G}{N}}\, dW_t^2 \\ &= -\frac{V_G}{V_S + V_P} z_t + \sqrt{\frac{V_G}{N}}\, dW_t^3, \end{aligned} \tag{S.3}$$

where $W_t^3$ is a standard Brownian motion. Eq. (S.3) confirms that, under stabilizing selection, the overall mean genetic value evolves according to an Ornstein–Uhlenbeck process (e.g., Lande 1976).

We now write $u_t = x_t/f$ and $v_t = y_t/(1-f)$. Then

$$du_t = \frac{dx_t}{f} = -\frac{V_G}{V_S + V_P} z_t + \sqrt{\frac{V_G}{lN}}\, dW_t^1, \tag{S.4}$$

$$dv_t = \frac{dy_t}{1-f} = -\frac{V_G}{V_S + V_P} z_t + \sqrt{\frac{V_G}{(1-f)N}}\, dW_t^2. \tag{S.5}$$

Define $w_t = u_t - v_t$. Taking the difference of Eqs. (S.4) and (S.5),

$$dw_t = du_t - dv_t = \sqrt{\frac{V_G}{lN}}\, dW_t^1 - \sqrt{\frac{V_G}{(1-f)N}}\, dW_t^2 = \sqrt{\frac{V_G}{f(1-f)N}}\, dW_t^4,$$

where $W_t^4$ is a standard Brownian motion. Therefore, $w_t = x_t/f - y_t/(1-f)$ behaves as a Brownian motion.

Under stabilizing selection, $z_t = x_t + y_t \approx 0$ for all $t$ large enough, and so, eventually, we can approximate $y_t = -x_t$. Therefore,

$$w_t = \frac{x_t}{f} - \frac{y_t}{1-f} = \frac{x_t}{f} + \frac{x_t}{1-f} = \frac{x_t}{f(1-f)},$$

and so

$$dx_t = f(1-f)dw_t = \sqrt{\frac{f(1-f)V_G}{N}}\, dW_t^4. \tag{S.6}$$

The chromosome-specific mean value $x_t$ therefore eventually behaves as a Brownian motion. With a starting value $x(0) = 0$, the mean of $x_t$ is zero and its variance is

$$\mathrm{Var}(x_t) = \frac{f(1-f)V_G}{N} \cdot t. \tag{S.7}$$

The formal justification for the approximation $z_t = x_t + y_t \approx 0$, and a complete characterization of the processes $x_t$and $y_t$, are given in Veller and Muralidhar (2025). It is shown that $x_t$ and $y_t$ tend to Brownian motions on a timescale of $V_S/V_G$ generations, the same timescale on which $z_t$ approaches its optimum, zero, having started away from the optimum.

Note that the rate of chromosome-specific drift is proportional to $V_G/N$, as also demonstrated via simulation in Fig. S5. Note furthermore that, in the absence of selection, the rate of chromosome-specific drift would be $fV_G/N$; the rate in Eq. (S.7) is reduced from this neutral rate by a factor $1-f$. This is the effect of selection on the trait. Thus, while chromosome-specific drift is qualitatively neutral in that it is Brownian, it is slower than neutral drift by an amount that depends on the contribution of the chromosome to the trait's genetic variance.

#### S1.2 Dynamics with sex chromosomes

The previous section illustrated the general principles of chromosome-specific drift. Here, I examine the case of chromosome-specific drift in a genome in which autosomes and sex chromosomes contribute to a polygenic trait under stabilizing selection. I consider a randomly mating species with male heterogamety (XX females and XY males) and of effective population size $N_e$. A quantitative trait is subject to stabilizing selection of (inverse) strength $V_S^{♂}$ in males and $V_S^{♀}$ in females. The phenotypic optimum is assumed to be the same in both sexes, although the dynamics are not affected in the long term by this assumption (Muralidhar and Coop 2024).

Let $x_t^{♂}$, $y_t^{♂}$, and $a_t^{♂}$ denote the mean haploid genetic contributions of the X, Y, and autosomes to the male trait in generation $t$, and let $x_t^{♀}$, $y_t^{♀}$, and $a_t^{♀}$ denote the analogous contributions to the female trait. Although females in the initial XX/XY system do not carry Y chromosomes, the Y nonetheless encodes a latent effect on the female phenotype that would be realized if some females came to carry Y chromosomes.

Given these haploid mean values of the various regions of the genome, the mean trait values in the initial male-heterogametic system are

$$z_t^{♂} = x_t^{♂} + y_t^{♂} + 2a_t^{♂}, \qquad z_t^{♀} = 2x_t^{♀} + 2a_t^{♀}.$$

We collect these into the vectors

$$\vec{x}_t = \begin{bmatrix} x_t^{♂} \\ x_t^{♀} \end{bmatrix}, \quad \vec{y}_t = \begin{bmatrix} y_t^{♂} \\ y_t^{♀} \end{bmatrix}, \quad \vec{a}_t = \begin{bmatrix} a_t^{♂} \\ a_t^{♀} \end{bmatrix}, \quad \vec{z}_t = \begin{bmatrix} z_t^{♂} \\ z_t^{♀} \end{bmatrix}.$$

Haploid genetic variances are denoted $V_G^{X♂}$, $V_G^{Y♂}$, $V_G^{A♂}$, $V_G^{X♀}$, $V_G^{Y♀}$, and $V_G^{A♀}$, and the haploid male-female genetic covariances of the various regions are denoted $C_G^X$, $C_G^Y$, and $C_G^A$. The total diploid genetic variances in each sex are $V_G^{♂}$ and $V_G^{♀}$, and the overall diploid male-female genetic covariance is $C_G$. These quantities are collected in the matrices

$$\mathbf{G}_X = \begin{pmatrix} V_G^{X♂} & C_G^X \\ C_G^X & V_G^{X♀} \end{pmatrix}, \quad \mathbf{G}_Y = \begin{pmatrix} V_G^{Y♂} & C_G^Y \\ C_G^Y & V_G^{Y♀} \end{pmatrix}, \quad \mathbf{G}_A = \begin{pmatrix} V_G^{A♂} & C_G^A \\ C_G^A & V_G^{A♀} \end{pmatrix}, \quad \mathbf{G} = \begin{pmatrix} V_G^{♂} & C_G \\ C_G & V_G^{♀} \end{pmatrix}.$$

In the absence of linkage disequilibrium, in the initial male-heterogametic system,

$$V_G^{♂} = V_G^{X♂} + V_G^{Y♂} + 2V_G^{A♂}, \qquad V_G^{♀} = 2V_G^{X♀} + 2V_G^{A♀}.$$

We assume that the X and Y chromosome do not recombine in males; any recombining portion of these chromosomes can be treated as autosomal. We further assume that the Y is nondegenerate, such that it contributes genetic variance comparable to the X, and that there is no dosage compensation in males, so that the effect of the Y in males is its haploid value.

Under these assumptions, the dynamics of the mean genetic values of the various regions for the male and female trait are described by the system of stochastic differential equations (SDEs):

$$\begin{aligned}
d\vec{x}_t &= -\begin{pmatrix} \frac{1}{3V_S^{♂}} & \frac{2}{3V_S^{♀}} \\ \frac{1}{3V_S^{♂}} & \frac{2}{3V_S^{♀}} \end{pmatrix} \odot \mathbf{G}_X \vec{z}_t \, dt + \sqrt{\tfrac{2}{3N_e}}\, \mathbf{G}_X^{1/2} d\vec{W}_t^X \\
d\vec{y}_t &= -\begin{pmatrix} \frac{1}{V_S^{♂}} & 0 \\ \frac{1}{V_S^{♂}} & 0 \end{pmatrix} \odot \mathbf{G}_Y \vec{z}_t \, dt + \sqrt{\tfrac{2}{N_e}}\, \mathbf{G}_Y^{1/2} d\vec{W}_t^Y \qquad \text{(S.8)} \\
d\vec{a}_t &= -\begin{pmatrix} \frac{1}{2V_S^{♂}} & \frac{1}{2V_S^{♀}} \\ \frac{1}{2V_S^{♂}} & \frac{1}{2V_S^{♀}} \end{pmatrix} \odot \mathbf{G}_A \vec{z}_t \, dt + \sqrt{\tfrac{1}{2N_e}}\, \mathbf{G}_A^{1/2} d\vec{W}_t^A,
\end{aligned}$$

where $\odot$ denotes element-wise multiplication and $\vec{W}_t^X$, $\vec{W}_t^Y$, and $\vec{W}_t^A$ are independent two-dimensional standard Brownian motions (Veller and Muralidhar 2025).

In Veller and Muralidhar (2025), we show these dynamics to lead to Brownian-like behavior of the mean contributions of the various regions to the male and female trait, despite stabilizing selection on their combined sums in both sexes, which instead behave like Ornstein–Uhlenbeck processes. Thus, over time, the X and Y chromosomes drift apart in their genetic values. This divergence is continually compensated by co-evolution among the chromosomes, maintaining the overall male and female mean genetic values at their optima.

When novel sexual genotypes arise, however, they disrupt this balance: newly rearranged sex chromosome genotypes are no longer co-adapted and lead to phenotypic displacement from the optimum. Because this chromosomal divergence behaves like a Brownian motion (see Fig. 1, S1), the degree of mismatch—and the associated fitness costs—increases steadily with the age of the resident sex chromosome system.

### S2 Quantifying the strength of resistance to heterogametic transitions

#### S2.1 Heterologous transitions in heterogamety

A heterologous transition in heterogamety via a new, dominant sex-determining mutation (here denoted $A$) involves six sexual genotypes (note that a heterologous transition in heterogamety involving a new recessive sex-determining mutation will involve the same six genotypes). If the transition occurs successfully, the population evolves from an initial state of entirely $aa\ XX$ females and $aa\ XY$ males to entirely $Aa\ YY$ females and $aa\ YY$ males; the latter state is one in which the former sex chromosome pair will behave autosomally, while the former autosomal chromosome pair serves as $W$ and $Z$ sex chromosomes.

Because this transition prompts the creation of females with $XY$ and $YY$ genotypes, and males with a $YY$ genotype, it may be impeded by fitness costs accruing to any of these novel sexual genotypes. In this model, these costs arise because these new sexual genotypes are not co-adapted, and therefore compensated, in their additive genetic values by the rest of the genome. Individuals with these genotypes may therefore be displaced off the phenotypic optimum and suffer reduced viability fitness.

All six genotypes and their corresponding viability fitnesses are given in the table below.

| | genotype | frequency | relative viability |
|---|---|---|---|
| ♀ | aa XX | $p_1$ | 1 |
| | Aa XX | $p_2$ | 1 |
| | Aa XY | $p_3$ | $1-r$ |
| | Aa YY | $p_4$ | $1-s$ |
| ♂ | aa XY | $p_5$ | 1 |
| | aa YY | $p_6$ | $1-t$ |

Sexual genotypes that were not present in the ancestral system—here, XY females and YY females and males—suffer fitness disadvantages under stabilizing selection, per the argument advanced in the Main Text. These fitness disadvantages are denoted $r$, $s$, and $t$ as in the table above.

Given some initial frequencies of the six genotypes, if the fitness disadvantages above are not too large, the frequencies converge rapidly to a 'slow manifold' along which the sex ratio is within a negligible distance of 1/2 (Bull and Charnov 1977; Veller et al. 2017). Label the frequencies of the genotypes $p_1, p_2, \ldots, p_6$ as in the table above, and assume that these frequencies lie on the slow manifold so that, to leading order, $p_1 + p_2 + p_3 + p_4 = 1/2$ and $p_5 + p_6 = 1/2$. Since the frequencies will stay in the close vicinity of the slow manifold, we need only derive update equations for $p_1$, $p_2$, $p_3$, and $p_5$.

Define

$$w_{♀} = \tfrac{1}{2} - p_3 r - p_4 s \tag{S.9}$$

$$= \frac{1}{2} - p_3 r - \left(\frac{1}{2} - p_1 - p_2 - p_3\right) s; \tag{S.10}$$

$$w_{♂} = \tfrac{1}{2} - p_6 t \tag{S.11}$$

$$= \frac{1}{2} - \left(\frac{1}{2} - p_5\right) t. \tag{S.12}$$

Then, after viability selection, the fractions of females with genotypes 1, 2, 3, and 4 are $v_1 = p_1/w_{♀}$, $v_2 = p_2/w_{♀}$, $v_3 = p_3(1-r)/w_{♀}$, and $v_4 = p_4(1-s)/w_{♀} = \left(\frac{1}{2} - p_1 - p_2 - p_3\right)(1-s)/w_{♀}$, while the fractions of males with genotypes 5 and 6 are $v_5 = p_5/w_{♂}$ and $v_6 = p_6(1-t)/w_{♂} = \left(\frac{1}{2} - p_5\right)(1-t)/w_{♂}$. Under random mating, the expected frequencies of genotypes 1, 2, 3, and 5 in the next generation are then

$$\begin{aligned}
p_1' &= \frac{1}{2}v_1v_5 + \frac{1}{4}v_2v_5 + \frac{1}{8}v_3v_5 \\
&=: f_1(p_1, p_2, p_3, p_5); \\
p_2' &= \frac{1}{4}v_2v_5 + \frac{1}{8}v_3v_5 \\
&=: f_2(p_1, p_2, p_3, p_5); \\
p_3' &= \frac{1}{4}v_2v_5 + \frac{1}{2}v_2v_6 + \frac{1}{4}v_3v_5 + \frac{1}{4}v_3v_6 + \frac{1}{4}v_4v_5 \\
&=: f_3(p_1, p_2, p_3, p_5); \\
p_5' &= \frac{1}{2}v_1v_5 + v_1v_6 + \frac{1}{4}v_2v_5 + \frac{1}{2}v_2v_6 + \frac{1}{4}v_3v_5 + \frac{1}{4}v_3v_6 + \frac{1}{4}v_4v_5 \\
&=: f_5(p_1, p_2, p_3, p_5).
\end{aligned} \tag{S.13}$$

We are interested in the potential for invasion of the initial aa XX, aa XY system, which corresponds to the equilibrium point $p^* = (p_1^*, p_2^*, p_3^*, p_5^*) = (1/2, 0, 0, 1/2)$. The Jacobian matrix of the system, evaluated at this initial point $p^*$, can be calculated to be

$$J|_{p^*} = \left.\begin{pmatrix} \frac{\partial f_1}{\partial p_1} & \frac{\partial f_1}{\partial p_2} & \frac{\partial f_1}{\partial p_3} & \frac{\partial f_1}{\partial p_4} \\ \frac{\partial f_2}{\partial p_1} & \frac{\partial f_2}{\partial p_2} & \frac{\partial f_2}{\partial p_3} & \frac{\partial f_2}{\partial p_4} \\ \frac{\partial f_3}{\partial p_1} & \frac{\partial f_3}{\partial p_2} & \frac{\partial f_3}{\partial p_3} & \frac{\partial f_3}{\partial p_4} \\ \frac{\partial f_4}{\partial p_1} & \frac{\partial f_4}{\partial p_2} & \frac{\partial f_5}{\partial p_3} & \frac{\partial f_4}{\partial p_4} \end{pmatrix}\right|_{p^*} = \begin{pmatrix} 1-s & \frac{1}{2}-s & \frac{1}{4}-s+\frac{3}{4}r & 1-t \\ 0 & \frac{1}{2} & \frac{1}{4}(1-r) & 0 \\ -\frac{1}{2}(1-s) & \frac{1}{2}s & \frac{1}{2}(s-r) & 0 \\ \frac{1}{2}(1-s) & -\frac{1}{2}s & \frac{1}{2}(r-s) & -(1-t) \end{pmatrix}. \tag{S.14}$$

We are interested in the eigenvalue of this Jacobian matrix that corresponds to the eigenvector $\sim (+0.853; -0.213; -0.426; +0.213)$, which is the eigenvector pointing in the direction of the slow manifold along which invasion of the system would proceed (the vector $(+0.853; -0.213; -0.426; +0.213)$ is the relevant eigenvector in the neutral system, and corresponds to an eigenvalue of 1 in that system).

#### S2.2 Homologous transitions in heterogamety

From an initial system of XX, XY male heterogamety, a mutated X chromosome X′ appears that is feminizing and, in this effect, is epistatically dominant to the Y chromosome. If X′ increases sufficiently in frequency relative to the wild-type X, the system transitions to one in which females have the genotype X′Y and males have the genotype YY, i.e., a system of female heterogamety where X′ serves as a W chromosome and Y serves as a Z chromosome. The transition between the old XX, XY system and the new X′Y, YY system involves intermediate states with five sexual genotypes, three female and two male:

| | genotype | frequency | relative viability |
|---|---|---|---|
| ♀ | XX | $p_1$ | 1 |
| | X′X | $p_2$ | 1 |
| | X′Y | $p_3$ | $1-s$ |
| ♂ | XY | $p_4$ | 1 |
| | YY | $p_5$ | $1-t$ |

Sexual genotypes that were not present in the ancestral system—here, X′Y females and YY males—suffer fitness disadvantages under stabilizing selection. These fitness disadvantages are denoted $s$ and $t$ respectively in the table above.

Given some initial frequencies of the five genotypes, if the fitness disadvantages above are not too large, then just as in the heterologous system above, the frequencies converge rapidly to a slow manifold along which the sex ratio is within a negligible distance of 1/2 (Bull and Charnov 1977; Veller et al. 2017). Label the frequencies of the genotypes $p_1, p_2, \ldots, p_5$ as in the table above, and assume that these frequencies lie on the slow manifold so that, to leading order, $p_1+p_2+p_3 = 1/2$ and $p_4+p_5 = 1/2$. Since the frequencies will stay in the close vicinity of the slow manifold, we need only derive update equations for $p_1$, $p_2$, and $p_4$.

Define

$$\begin{aligned} w_{♀} &= \tfrac{1}{2} - p_3 s && \text{(S.15)} \\ &= \frac{1}{2} - \left(\frac{1}{2} - p_1 - p_2\right) s; && \text{(S.16)} \\ w_{♂} &= \tfrac{1}{2} - p_5 t && \text{(S.17)} \\ &= \frac{1}{2} - \left(\frac{1}{2} - p_4\right) t. && \text{(S.18)} \end{aligned}$$

Then, after viability selection, the fractions of females with genotypes 1, 2, and 3 are $v_1 = p_1/w_{♀}$ and $v_2 = p_2/w_{♀}$, and $v_3 = p_3(1-s)/w_{♀} = \left(\frac{1}{2} - p_1 - p_2\right)(1-s)/w_{♀}$, while the fractions of males with genotypes 4 and 5 are $v_4 = p_5/w_{♂}$ and $v_5 = p_5(1-t)/w_{♂} = \left(\frac{1}{2} - p_4\right)(1-t)/w_{♂}$. Under random mating, the expected frequencies of genotypes 1, 2, and 4 in the next generation are then

$$\begin{aligned} p_1' &= \frac{1}{2}v_1 v_4 + \frac{1}{4}v_2 v_4 \\ &=: f_1(p_1, p_2, p_4); \\ p_2' &= \frac{1}{4}v_2 v_4 + \frac{1}{4}v_3 v_4 \\ &=: f_2(p_1, p_2, p_3, p_5); \\ p_4' &= \frac{1}{2}v_1 v_4 + v_1 v_5 + \frac{1}{4}v_2 v_4 + \frac{1}{2}v_2 v_5 + \frac{1}{4}v_3 v_4 \\ &=: f_4(p_1, p_2, p_3, p_5). && \text{(S.19)} \end{aligned}$$

We are interested in the potential for invasion of the initial XX, XY system, which corresponds to the equilibrium point $p^* = (p_1^*, p_2^*, p_4^*) = (1/2, 0, 1/2)$. The Jacobian matrix of the system, evaluated at this initial point $p^*$, can be calculated to be

$$J|_{p^*} = \left( \begin{array}{ccc} \frac{\partial f_1}{\partial p_1} & \frac{\partial f_1}{\partial p_2} & \frac{\partial f_1}{\partial p_4} \\ \frac{\partial f_2}{\partial p_1} & \frac{\partial f_2}{\partial p_2} & \frac{\partial f_2}{\partial p_4} \\ \frac{\partial f_4}{\partial p_1} & \frac{\partial f_3}{\partial p_2} & \frac{\partial f_4}{\partial p_4} \end{array} \right) \Bigg|_{p^*} = \left( \begin{array}{ccc} 1-s & \frac{1}{2}-s & 1-t \\ -\frac{1}{2}(1-s) & \frac{1}{2}s & 0 \\ \frac{1}{2}(1-s) & -\frac{1}{2}s & -(1-t) \end{array} \right). \tag{S.20}$$

We are interested in the eigenvalue of this Jacobian matrix that corresponds to the eigenvector $\sim (+0.873; -0.436; +0.218)$, which is the eigenvector pointing in the direction of the slow manifold along which invasion of the system would proceed (the vector $(+0.873; -0.436; +0.218)$ is the relevant eigenvector in the neutral system, and corresponds to an eigenvalue of 1 in that system).

#### S2.3 The eigenvalues predict the probability of transition and can thus be thought of as selection coefficients

To show that the eigenvalues described above predict the fixation probability of a new sex-determining mutation and thus transition of the sex-determining system, I estimate these transition probabilities using Monte-Carlo simulations for various configurations of the selection coefficients on the novel sexual genotypes produced in the course of invasion of the new sex-determining mutation, both for the heterologous system and the homologous system, and concentrating on epistatically dominant new sex-determining mutations ($A$ in the heterologous system; $X'$ in the homologous system).

For each configuration, I randomly sampled a maximum fitness effect $s_{\max}$ from the interval $[-0.01, 0.01]$ (half of the time from a uniform distribution on the interval, and half of the time from a normal distribution with mean 0 and standard deviation 0.01/2, discarding draws outside of $[-0.01, 0.01]$). I then chose the selection coefficients of the novel sexual genotypes ($r$, $s$, and $t$ for a heterologous transition; $s$ and $t$ for a homologous transition) independently from a uniform distribution on $[0, s_{\max}]$ if $s_{\max} > 0$ or $[s_{\max}, 0]$ if $s_{\max} < 0$. Note that this sampling scheme guarantees that the novel sexual genotypes are either all deleterious or all beneficial; this ensures that there are no stable internal equilibria with all genotypes present (Orzack et al. 1980), which would potentially hang the simulations described below.

For each configuration of selection coefficients, the eigenvalues and eigenvectors of the Jacobian matrix (Eq. S.14 or S.20) were computed. The relevant eigenvalue $\lambda_{\text{inv}}$ was chosen as that corresponding to the eigenvector closest (in Euclidean distance) to $(+0.853; -0.213; -0.426; +0.213)$ for the heterologous case and $(+0.873; -0.436; +0.218)$ for the homologous case. The effective selection coefficient on the new sex-determining mutation was taken to be $s_{\text{eff}} = \lambda_{\text{inv}} - 1$.

Having chosen the selection coefficients for a particular configuration and computed $s_{\text{eff}}$, I then simulated $10^7$ Wright-Fisher trials with population size $N = 1{,}000$. For the heterologous case, each trial started with a single $A$ mutation, half the time in an $AaXX$ female and half the time in an $AaXY$ female. For the homologous case, each trial started with a single $X'$ mutation, half the time in an $X'X$ female and half the time in an $X'Y$ female. The starting numbers of males and females were $N/2$ in both cases. Each trial was run until only the two initial sexual genotypes were present ($aaXX$ and $aaXY$ in the heterologous case; $XX$ and $XY$ in the homologous case) or the two sexual genotypes constituting the new heterogametic system were present ($AaYY$ and $aaYY$ in the heterologous case; $X'Y$ and $YY$ in the homologous case); the latter trials were scored as fixations of the new sex-determining mutation. The transition probability was estimated as the proportion of trials scored as fixations.

Figure S15 plots the values of $s_{\text{eff}}$ against these estimated transition probabilities for the cases of heterologous (Fig. S15A,B) and homologous (Fig. S15C,D) transitions. Three observations are of note. First, for both cases, the transition probability is a nearly monotonic function of $s_{\text{eff}}$; in this sense, $s_{\text{eff}}$ (and thus the relevant eigenvalue $\lambda_{\text{inv}}$) is an effective predictor of the transition probability and thus an effective summary of the selection coefficients on the novel sexual genotypes in a particular configuration. Second, outside of the nearly-neutral regime ($N|s_{\text{eff}}| > 1$ ), the fixation probability corresponds approximately to the classical formula for fixation of a mutant allele in a population of size $N$ (Kimura 1962), substituting $s_{\text{eff}}$ for the selection coefficient of the allele (Fig. S15A,C). This close correspondence justifies the use of the term 'effective selection coefficient'. Third, inside the nearly neutral regime $N|s_{\text{eff}}| < 1$, the fixation probability is higher than predicted by the classical fixation probability formula (Fig. S15B,D). This is the result of 'drift-induced selection' in favor of epistatically dominant sex-determining mutations in both heterologous and homologous transitions (Veller et al. 2017).

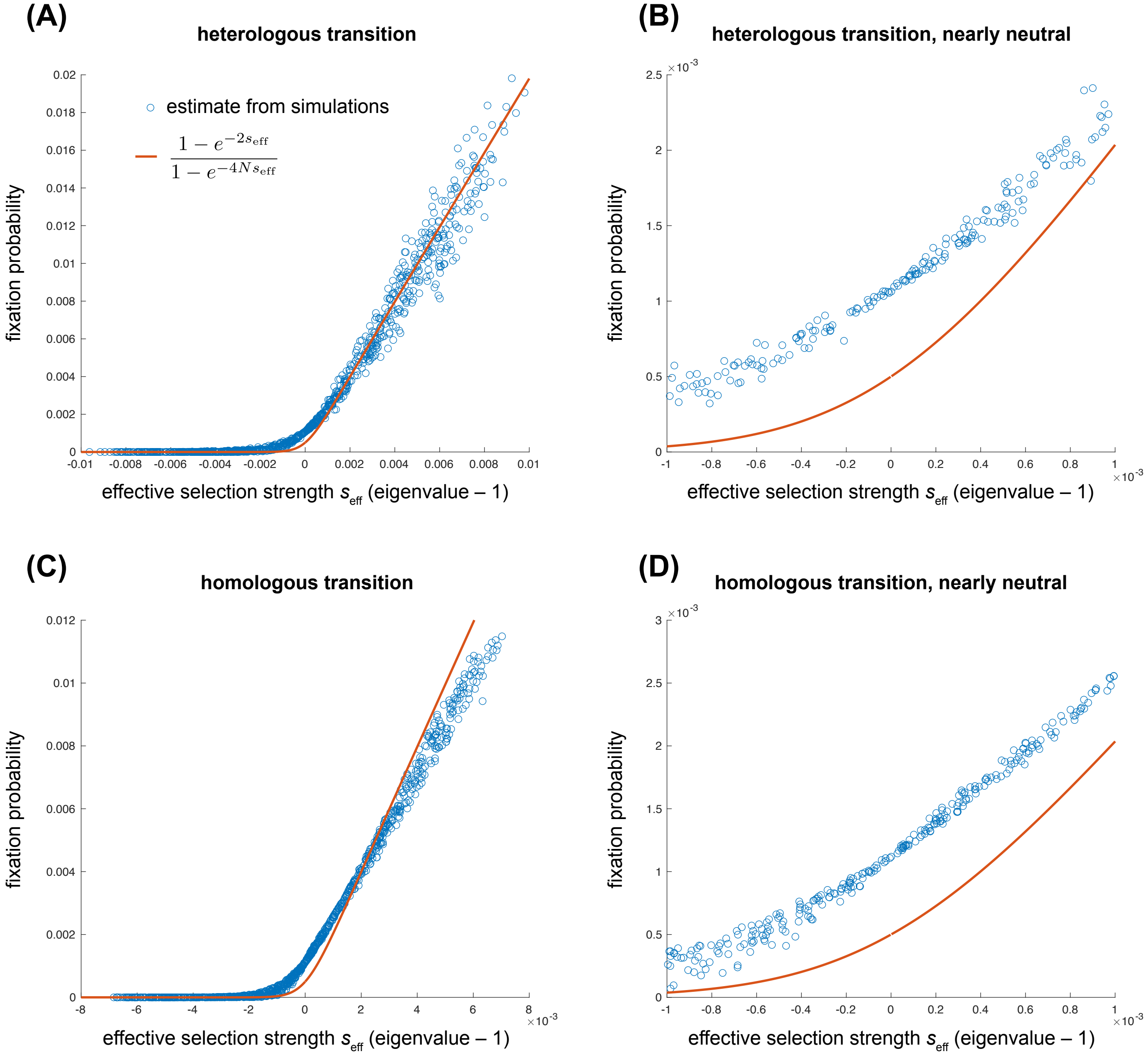


**Figure S15:** Effective selection coefficients (invasion eigenvalues minus one) for various configurations of the fitness costs of novel sexual genotypes (x axis), and the associated probabilities of sex chromosome turnover estimated via Monte Carlo simulation (y axis), for heterologous (panels A and B) and homologous (panels C and D) turnovers of the system of heterogamety seeded by a single copy of a new, epistatically dominant major sex-determining mutation. The Monte Carlo simulations are described in the Methods and in Section S2.3 above. Red lines show the fixation probability formula of Kimura (1962). Panels B and D zoom in on the nearly-neutral regime ($|Ns_{\rm eff}| < 1$), where the empirical transition probabilities are higher than predicted by Kimura's formula because of the phenomenon of drift-induced selection in favor of dominant major sex-determining alleles (Veller et al. 2017).
